# Epitenon-derived cells comprise a distinct progenitor population that contributes to both fibrosis and regeneration following acute flexor tendon injury in a spatially-dependent manner

**DOI:** 10.1101/2023.01.30.526242

**Authors:** Anne E.C. Nichols, Lauren Benoodt, Emmanuella Adjei-Sowah, Kyle Jerreld, Alexander Kollar, Constantinos Ketonis, Alayna E. Loiselle

**Author notes:** **Corresponding Author:** Anne E.C. Nichols, PhD, Center for Musculoskeletal Research, University of Rochester Medical Center, 601 Elmwood Ave, Box 665G, Rochester, NY 14642.

## Abstract

Flexor tendon injuries are common and heal poorly owing to both the deposition of function-limiting peritendinous scar tissue and insufficient healing of the tendon itself. Therapeutic options are limited due to a lack of understanding of the cell populations that contribute to these processes. Here, we identified the epitenon as a major source of cells that contribute to both peritendinous fibrosis and regenerative tendon healing following acute tendon injury. Using a combination of genetic lineage tracing and single cell RNA-sequencing (scRNA-seq), we profiled the behavior and contributions of each cell fate to the healing process in a spatio-temporal manner. Integrated scRNA-seq analysis of mouse healing with human peritendinous scar tissue revealed remarkable transcriptional similarity between mouse epitenon-derived cells and fibroblasts present in human peritendinous scar tissue, which was further validated by immunofluorescent staining for conserved markers. Finally, ablation of pro-fibrotic epitenon-derived cells post-tendon injury significantly improved functional recovery. Combined, these results clearly identify the epitenon as the cellular origin of an important progenitor cell population that could be leveraged to improve tendon healing.

## INTRODUCTION

Tendons play an essential role in musculoskeletal movement as the means by which muscle contractions are transmitted to bone. Acute tendon injuries occur frequently and heal poorly due to two separate but inextricably linked issues: 1) an exuberant fibrotic healing process that reduces tendon range of motion (ROM) via the formation of peritendinous fibrovascular scar tissue around the injury site and 2) poor intrinsic healing capacity resulting in sub-optimal regeneration of native tendon structure^1^. Injuries to the flexor tendons of the hand are particularly common (more than 300,000 per year^2^) and generate an enormous socioeconomic burden estimated at greater than $400 million per year in the United States^3^. Beyond an initial surgery to restore tendon continuity, up to 24%^4–6^ of patients with injured flexor tendons require a second surgical procedure (tenolysis) to remove peritendinous scar tissue and restore ROM, further prolonging patient recovery and increasing the risk of rupture. Despite this significant clinical problem, the ability to design targeted therapeutics to improve tendon healing is severely limited due to a lack of understanding of how various cell populations contribute to the healing process. In particular, the specific identity, origins, and molecular programming of the cells that deposit and sustain the peritendinous scar tissue remain unknown.

Initial work describing flexor tendon healing in canine and non-human primate models identified the epitenon (a thin layer of cells on the surface of tendons) as the primary orchestrator of the early tendon healing response, with epitenon-derived cells thought to contribute to the deposition of collagen in and around the injury site^7–10^ and to the production of pro-fibrotic growth factors and cytokines^11^ that could potentially contribute to fibrovascular scar formation. Unfortunately, lack of epitenon-specific markers to date has precluded further study of this important cell population. Given the important speculated role of the epitenon as a contributor to the formation of peritendinous adhesions, we sought to identify a genetic driver that could be used to target, track, and manipulate epitenon cells. In particular, we focused on a BAC transgenic mouse line that encodes a tamoxifen (TMX)-inducible Cre (*Cre^ERT^*) under the control of the *GLAST* [glial high affinity glutamate transporter] (also known as *Slc1a3* or *EAAT1*) promoter^12^. The *GLASTCre^ERT^* driver has previously been used to target various types of mesenchymal progenitor cells in other tissues including muscle^13, 14^ and the spinal cord^15^ that ultimately contribute to both the physiological healing process and pathological processes such as fibrosis^15^ and heterotopic ossification^13, 14^. This particular driver was therefore an attractive candidate for labeling a population with similar purported function in tendon.

In the present study, we identified the *GLASTCre^ERT^*driver and gene product (GLAST) as a specific marker of epitenon cells in adult mouse and human flexor tendons. We then utilized this new genetic tool to trace the behavior and fate of epitenon-derived cells following acute tendon injury using a combination of genetic lineage tracing and single-cell RNA-sequencing (scRNA-seq) in mice. Surprisingly, epitenon-derived cells were found to contribute not only to fibrotic tendon healing, but also to tendon regeneration in a spatially-dependent manner. Ablation of pro-fibrotic epitenon-derived cells post-injury significantly improved functional recovery. Finally, pro-fibrotic epitenon-derived mouse cells were found to be transcriptionally similar to those found in human peritendinous scar tissue, implicating epitenon cells in the formation of function-limiting adhesive scar in human flexor tendon injuries and identifying them as a potential therapeutic target for future study. Combined, these data provide the first in-depth characterization of the epitenon cellular environment and identify the epitenon as a niche for a bi-fated progenitor cell population that contributes to both tendon regeneration and fibrosis following acute injury.

## RESULTS

### Characterization of the epitenon cellular niche during tendon homeostasis

To determine the targeting of the *GLASTCre^ERT^* driver during tendon homeostasis, mice were crossed to the *ROSA-Ai9* reporter strain (*GLASTCre^ERT^; Ai9^F/F^*, referred to as GLAST^Ai9^ mice) which contains the transcript encoding the red fluorescent protein tdTomato downstream of a floxed stop cassette^16^ (**Fig. 1A**). Upon tamoxifen (TMX) administration, Cre-mediated recombination causes removal of the stop cassette, allowing permanent expression of tdTomato in all cells expressing *GLAST*, thus labeling them, and all progeny, as GLAST^Ai9^ cells. Intact flexor digitorum longus (FDL) tendons were examined 14 days after TMX treatment using several imaging modalities (**Fig. 1A**). Sagittal sections through the hind paw demonstrate that GLAST^Ai9^ cells were largely found in a single-cell thick layer lining the surface of both the superficial and deep FDL tendons (**Fig. 1B**), consistent with prior characterization of the epitenon structure^17^. A small number of GLAST^Ai9^ cells were also observed within the tendon body (**Fig. 1B**, **1C**). Similarly, GLAST^Ai9^ cells were found lining other tendons including the flexor carpi ulnaris tendon in the forelimb, the tendons in the tail, and the Achilles tendon (**Fig. S1A**). In contrast, very few GLAST^Ai9^ cells were found in the patellar tendon epitenon (**Fig. S1B**). Previous reports have indicated a small percentage of non-specific recombination using this driver^12, 18^; however, no cells were labeled in the epitenon or in the tendon body without exposure to TMX (**Fig. S1C**).

**Fig. 1.**
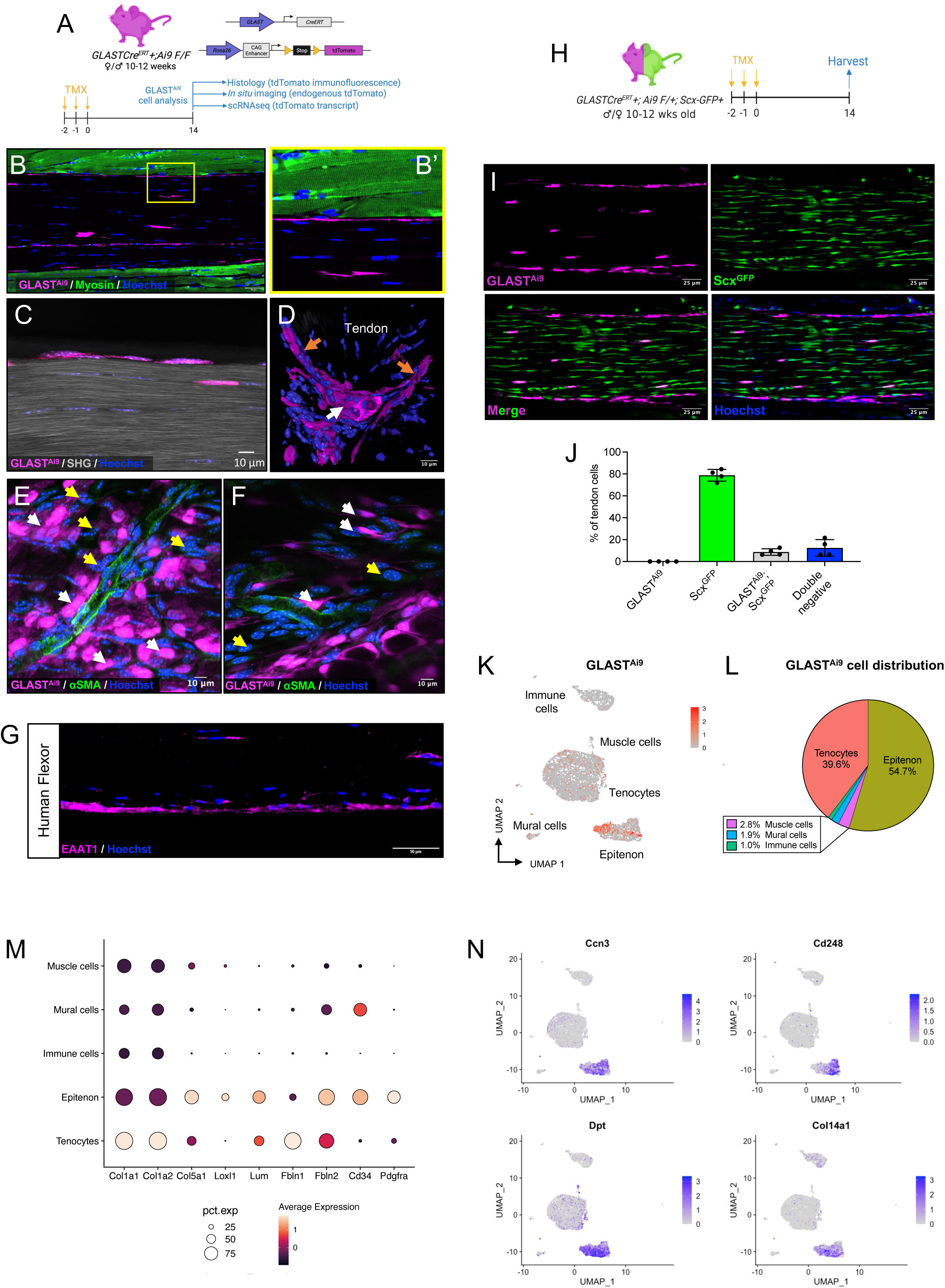
Characterization of the tendon epitenon during homeostasis. (**A**) Overview of experimental design. (**B**) Sagittal section of the FDL tendon showing IF for GLAST^Ai9^ (magenta) and myosin (green). Scale bar = 50 μm. (**C**) SHG imaging (gray) of endogenous GLAST^Ai9^ labeling (magenta) in relation to the collage matrix in sagittal frozen sections. (**D**) 3D reconstruction of an axial section through the FDL tendon showing GLAST^Ai9^ (magenta) cells in both the epitenon (orange arrows) and surrounding blood vessels adjacent to the epitenon (white arrows). (**E**) In situ multiphoton imaging of GLAST^Ai9^ epitenon cells (magenta cells; white arrows), small blood vessels on the surface of the FDL tendon (αSMA; green), and non-GLAST^Ai9^ epitenon cells (yellow arrows). (**F**) In situ multiphoton imaging of GLAST^Ai9^ perivascular cells (white arrows) and non-GLAST^Ai9^ cells (yellow arrows). Scale bars = 10 μm. (**G**) Representative immunofluorescent staining for EAAT1 (magenta) protein encoded by GLAST gene) in human flexor tendon. Scale bar = 50 μm. (**H**) GLAST^Ai9^; Scx^GFP^ mice received TMX injections and were harvested for frozen histology. (**I**) Sagittal section (scale bar = 25 μm) and (**J**) quantification of the mean endogenous GLAST^Ai9^ (magenta) and Scx^GFP^ (green) fluorescence. Error bars represent standard deviation. UMAP of clusters containing GLAST^Ai9^ cells (**K**) in scRNA-seq dataset from uninjured FDL tendons. (**L**) GLAST^Ai9^ cell distribution in cell clusters. (**M**) Expression of pan-fibroblast marker genes in clusters from uninjured tendon. (**N**) Transcriptional markers of the epitenon cell cluster in uninjured FDL tendons. All images are representative of n = 4-6 mice. In all images, nuclei are counterstained with Hoechst (blue).

To evaluate the localization of GLAST^Ai9^ cells relative to the collagenous matrix of the tendon, we utilized second harmonic generation (SHG) microscopy. Consistent with the idea that GLAST^Ai9^ cells are located in the epitenon on the surface of the tendon rather than within the tendon itself, GLAST^Ai9^ cells were located outside the tendon matrix (**Fig. 1C**) in the space between the tendon and the adjacent muscle. 3D reconstruction of axial tissue sections taken through the FDL tendon further revealed a distinct boundary between the GLAST^Ai9^ cells on the surface of the tendon and the tenocytes within (**Fig. 1D**). These data also indicate that the *GLASTCre^ERT^* driver targets multiple cell populations within the epitenon space, including perivascular cells. To further evaluate the cellular makeup of the epitenon niche, we developed an *in situ* multiphoton imaging protocol. Using this approach, we confirmed the presence of both perivascular and non-perivascular GLAST^Ai9^ cells (**Fig. 1E**, **1F**; white arrows) as well as populations of non-GLAST^Ai9^ cells throughout the epitenon (**Fig. 1E**, **1F**; yellow arrows). Immunofluorescent staining for excitatory amino acid transporter 1 (EAAT1; the protein encoded by *Slc1a3*/GLAST) in healthy human flexor tendon (**Fig. 1G**) demonstrated that GLAST can also be used to mark the epitenon in human flexor tendons. In addition to the epitenon, 9.1 ± 4.9% of human tenocytes were EAAT1+.

Due to the presence of a small number of GLAST^Ai9^ cells located within the tendon body (**Fig. 1B**, **1C**) we next sought to evaluate the relationship between GLAST^Ai9^ cells and scleraxis (Scx)-expressing tenocytes by generating dual reporter mice to track active *Scx* expression (Scx^GFP^)^19^ in addition to labeling GLAST^Ai9^ cells (**Fig. 1H**; GLAST^Ai9^; Scx^GFP^ mice). GLAST^Ai9^ cells located in the epitenon did not express Scx-GFP (**Fig. 1I**) indicating that this population is distinct from previously described adult Scx-lineage tendon cells^19, 20^. Similar to the human flexor tendon, a small number (8.7 ± 3%) of GLAST^Ai9^; Scx-GFP+ tenocytes (**Fig. 1J**) were also observed in the body of the mouse FDL tendon.

In order to better understand the identity of GLAST^Ai9^ cells during tendon homeostasis, we performed single cell RNA-sequencing (scRNA-seq) on uninjured FDL tendons from GLAST^Ai9^ mice. Initial low-resolution clustering revealed five main populations (**Fig. 1K, S1D**): tenocytes (*Scx+*), immune cells (*Adgre1+*), mural cells (*Pecam1+*), muscle cells (*Myod1+*), and a cluster that we subsequently defined as epitenon cells based on the relatively high percentage of GLAST^Ai9^+ cells in this cluster (**Fig. 1L, S1D**). Confirming our imaging observations, the majority of GLAST^Ai9^ cells (54.7%) were located within the epitenon cell cluster (23.7% of all epitenon cells), with smaller numbers found within the tenocyte (5.7% of all tenocytes; 39.6% of GLAST^Ai9^ cells), muscle (6.7% of all muscle cells; 2.8% of GLAST^Ai9^ cells), mural (7.1% of all mural cells; 1.9% of GLAST^Ai9^ cells), and immune (2.1% of all immune cells; 1.0% of GLAST^Ai9^ cells) cell clusters (**Fig. 1L**). Similar to our dual reporter (GLAST^Ai9^; Scx^GFP^) imaging, a small percentage (3.3%; **Fig. S1E**) of GLAST^Ai9^ tenocytes were found within the tenocyte cluster. Expression of pan-fibroblast markers^21^ including collagen types 1 alpha 1 and 2 (*Col1a1* and *Col1a2*), collagen type 5 alpha 1 (*Col5a1*), lysyl oxidase like 1 (*Loxl1*), lumican (*Lum*), *Cd34*, and platelet-derived growth factor alpha (*Pdgfra*) indicated that epitenon cells are a fibroblast population that is markedly distinct from tenocytes (**Fig. 1M**).

### Identification of epitenon-specific transcriptional markers

Given the historic lack of specific epitenon markers for transcriptomic studies, we next interrogated our scRNA-seq dataset to identify candidate epitenon markers that do not rely on lineage tracing. Expression of communication network factor 3 (*Ccn3*), endosialin (*Cd248*), dermatopontin (*Dpt*), and collagen type 14 alpha 1 (*Col14a1*), (**Fig. 1N)** were found to be specific transcriptional markers for the epitenon cluster. A table containing all epitenon cluster markers can be found in **Supplemental Table 1**.

### Epitenon cells proliferate and migrate to the scar area following acute tendon injury

Prior work speculated that epitenon-derived cells are early contributors to the tendon healing response; however, this has yet to be definitively demonstrated. To determine the extent to which epitenon-derived cells participate in tendon healing, GLAST^Ai9^; Scx^GFP^ mice underwent FDL tendon injury and repair surgery, and tendons were examined at days 3, 7, 10, 14, 21, and 28 post-injury (**Fig. 2A**). GLAST^Ai9^ cells were labeled prior to injury and included a washout period to ensure that only cells expressing GLAST prior to injury were labeled and traced. At day 3 (D3) post-injury, increased numbers of GLAST^Ai9^ cells were observed in the epitenon at the periphery of the injury site (**Fig. 2B**), with expansion of the epitenon through D7. Beginning at D10, GLAST^Ai9^ cells were found both within the newly formed, organized bridging tendon tissue that forms between the cut tendon ends as well as on the periphery of the injury site. This localization pattern persisted relatively unchanged through at least D28.

**Fig. 2.**
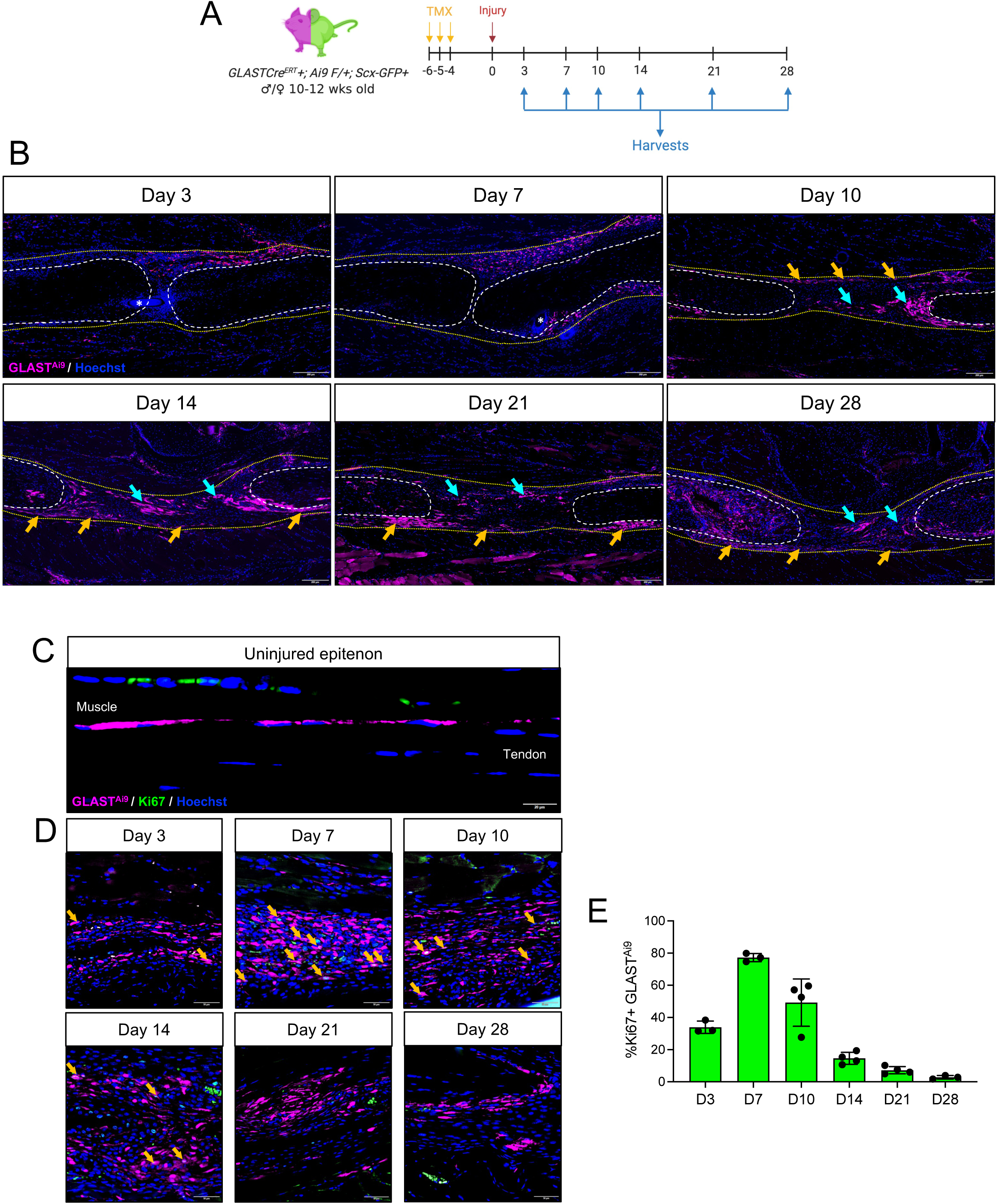
GLAST^Ai9^ epitenon cells proliferate and migrate into the scar area follow acute tendon injury. (**A**) Overview of experimental design. (**B**) Representative images of GLAST^Ai9^ (magenta) cells located in both a peripheral capsule (orange arrows) around the injury site and within the scar tissue (aqua arrows). Tendon stubs are outlined by white dashed lines and the scar area is outlined by yellow dotted lines. Sutures are marked by white asterisks. Scale bar = 200 μm. (**C**) Ki67 IF (green) in the uninjured epitenon (GLAST^Ai9^; magenta) and representative images (**D**) of the epitenon adjacent to the injury site throughout healing. (**E**) Quantification of the mean percentage of proliferating GLAST^Ai9^ epitenon cells (GLAST^Ai9^;Ki67+) at each timepoint. Error bars represent standard deviation. In all images, nuclei are counterstained with Hoechst (blue). All images are representative of n = 3-4 mice per timepoint.

Staining for the proliferation marker Ki67 indicated that GLAST^Ai9^ cells do not proliferate in the epitenon during homeostasis (**Fig. 2C**); however, shortly after injury (∼D3), Ki67+ GLAST^Ai9^ cells (34.0 ± 3.8%) were observed in the expanded epitenon adjacent to the injury site (**Fig. 2B**, **2C**). The number of Ki67+ GLAST^Ai9^ cells in the epitenon peaked at D7 (77.2 ± 2.6%) before declining through D28 (D10: 49.2 ± 14.7%, D14: 14.6 ± 3.7%, D21: 7.2 ± 2.3%, D28: 2.8 ± 1.1%).

### Epitenon-derived cells exhibit spatially-dependent fates

Given the spatially distinct localization of GLAST^Ai9^ epitenon-derived cells (peripheral capsule vs. organized bridging tissue), we next assessed the relationship between GLAST^Ai9^ cells and previously identified tenocyte populations (Scx-GFP+)^20, 22^ throughout healing. Prior to D10, GLAST^Ai9^ and Scx-GFP+ cells in and around the injury site remained largely distinct populations. Interestingly, GLAST^Ai9^; Scx-GFP+ cells began to appear in the organized bridging tissue at around D10 (**Fig. 3A**, **3B**; white arrows), indicating differentiation of GLAST^Ai9^ epitenon cells into tenocytes post-injury. The overall proportion of GLAST^Ai9^; Scx-GFP+ in the organized bridging tissue increased over time through D28 (**Fig. 3C**; D3: 1.7 ± 1.0%, D7: 3.5 ± 4.0%, D10: 5.9 ± 4.0%, D14: 2.5 ± 0.9%, D21: 10.1 ± 2.8%, D28: 15.3 ± 1.4%). At D28, approximately one-third (33.0 ± 6.7%) of the Scx-GFP+ tenocytes in the organized bridging tissue were GLAST^Ai9^ indicating that the epitenon serves as a major source of tenogenic progenitors following injury. Given the small number of GLAST^Ai9^; Scx-GFP+ cells observed within the tendon body prior to injury (**Fig. 1A**) and our previous work demonstrating that tenocytes within the tendon stubs adjacent to the injury site activate to myofibroblasts following injury^20, 23^, we stained for alpha smooth muscle actin (αSMA; a marker of mature myofibroblasts) to determine if GLAST^Ai9^; Scx-GFP+ cells follow a similar fate trajectory post-injury (**Fig. S3A**). Starting at D14, GLAST^Ai9^; αSMA+ cells were observed within the stubs suggesting that GLAST^Ai9^ tenocytes contribute to the post-injury myofibroblast population. Several tenogenic progenitor cell populations that contribute to tendon healing have previously been identified, including melanoma cell adhesion molecule (Mcam; CD146+) cells^24, 25^. Immunofluorescent staining for CD146 (**Fig. S3B**) revealed no GLAST^Ai9^; CD146+ cells at any point during tendon healing indicating that GLAST^Ai9^ epitenon cells represent a distinct progenitor pool from CD146+ cells.

**Fig. 3.**
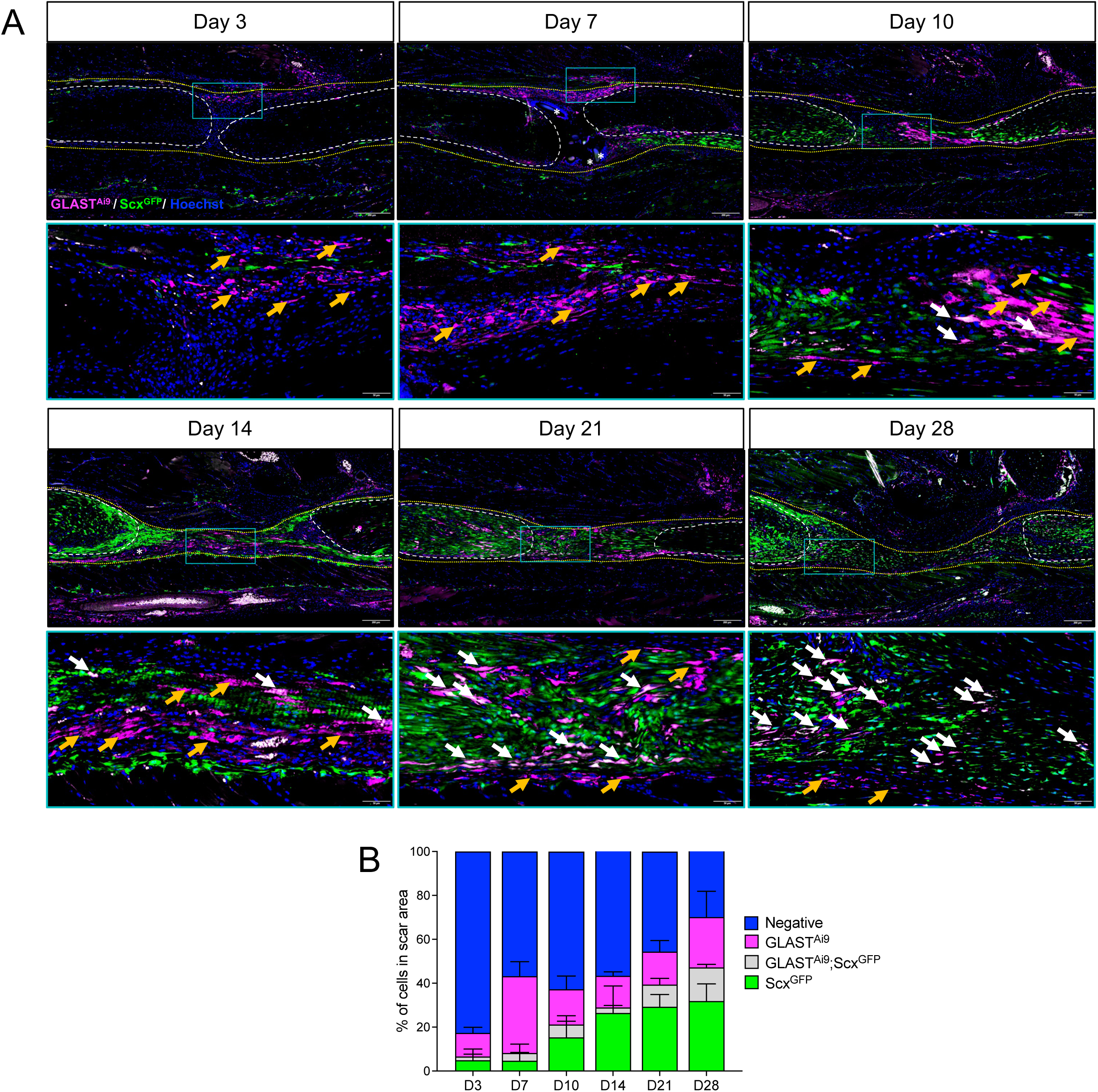
A subset of GLAST^Ai9^ epitenon cells differentiate into tenocytes during healing. (**A**) Representative images of healing tendons from GLAST^Ai9^; Scx^GFP^ mice at days 3, 7, 10, 14, 21, and 28 post-injury showing the presence of GLAST^Ai9^; Scx^GFP^-cells (magenta cells, orange arrows), Scx^GFP^+ tenocytes (green cells), and GLAST^Ai9^; Scx^GFP^+ tenocytes (white cells, white arrows). Tendon stubs are outlined by white dashed lines and the scar area is outlined by yellow dotted lines. Sutures are marked by white asterisks. Scale bars = 200 μm, magnified image scale bars = 50 μm. (**B**) Quantification of all cell populations present in the scar area throughout healing. Bars represent the mean cell number and error bars denote the standard deviation. In all images, nuclei are counterstained with Hoechst (blue). All images are representative of n = 3-4 mice per timepoint.

In addition to GLAST^Ai9^; Scx-GFP+ cells in the organized bridging tissue, we observed a persistent band of GLAST^Ai9^; Scx-GFP-cells in a peripheral capsule surrounding the injury site in the region primarily associated with the formation of fibrotic, peritendinous adhesions^23^ from D14 on (**Fig. 3A**; orange arrows). Combined, these lineage tracing experiments suggest two potential fates for epitenon cells post-injury: 1) tenogenic differentiation within the organized bridging tissue or 2) involvement in the formation of peritendinous fibrotic scar tissue.

### Single-cell transcriptomic analysis identifies temporally-dynamic contributions of epitenon-derived cells to tendon healing

To better understand the overall role of the epitenon during tendon healing, we performed scRNA-seq on healing tendons from GLAST^Ai9^ mice at 7, 14, and 28 days post-injury. These timepoints were chosen based on our lineage tracing experiments (**Figs. 2 and 3**) and represent points at which key transitional events appeared to occur in epitenon-derived cells: D7 – expansion of the epitenon (large numbers of Ki67+ GLAST^Ai9^ cells), D14 – migration into the fibrotic capsule (GLAST^Ai9^; Scx-GFP-cells) or tenogenic differentiation in the organized bridging tissue (few GLAST^Ai9^; Scx-GFP+ cells), and D28 – persistence in the fibrotic capsule (GLAST^Ai9^; Scx-GFP-cells) or maturation into tenocytes (increased numbers of GLAST^Ai9^; Scx-GFP+ cells in the organized bridging tissue). To facilitate tracing of the epitenon cells throughout healing, we first integrated the post-injury timepoints with the uninjured dataset described above (**Fig. 4A**) and used the GLAST^Ai9^ genetic label to identify epitenon-derived cells in the integrated dataset (**Fig. 4B**). The integrated dataset captured a wide variety of cell types including several macrophage populations, pericytes, injury-associated fibroblasts (IAF), mural cells, T-cells, and neutrophils which were annotated using canonical markers (**Fig. 4B**). This comprehensive scRNA-seq atlas of tendon healing will be described in detail elsewhere. For the purpose of this study, we focused our investigation on the clusters containing the largest proportion of epitenon-derived cells (GLAST^Ai9^). In addition to the epitenon cell cluster (**Fig. 4C**), an increasing proportion of GLAST^Ai9^ cells (uninjured: 11.9%, D7: 27.3%, D14: 24.7%, D28: 23.2%) was also found in a second cluster that was largely absent in the uninjured tendon but expanded dramatically post-injury (**Fig. 4C**). We surmised that this population most likely represents the bridging scar tissue given the presence and increasing proportion of GLAST^Ai9^; Scx-GFP+ cells in this cluster (**Fig. 4C**) and as seen by IF (**Fig. 3**). Based on these observations, we subsequently defined this cluster as the bridging cell cluster. As there were no significantly differentially expressed genes (DEG) between GLAST^Ai9+^ and GLAST^Ai9-^ cells in either the epitenon or bridging cell clusters (**Fig. S4A**), the fate of each population was examined independent of GLAST lineage for all subsequent analyses.

**Fig. 4.**
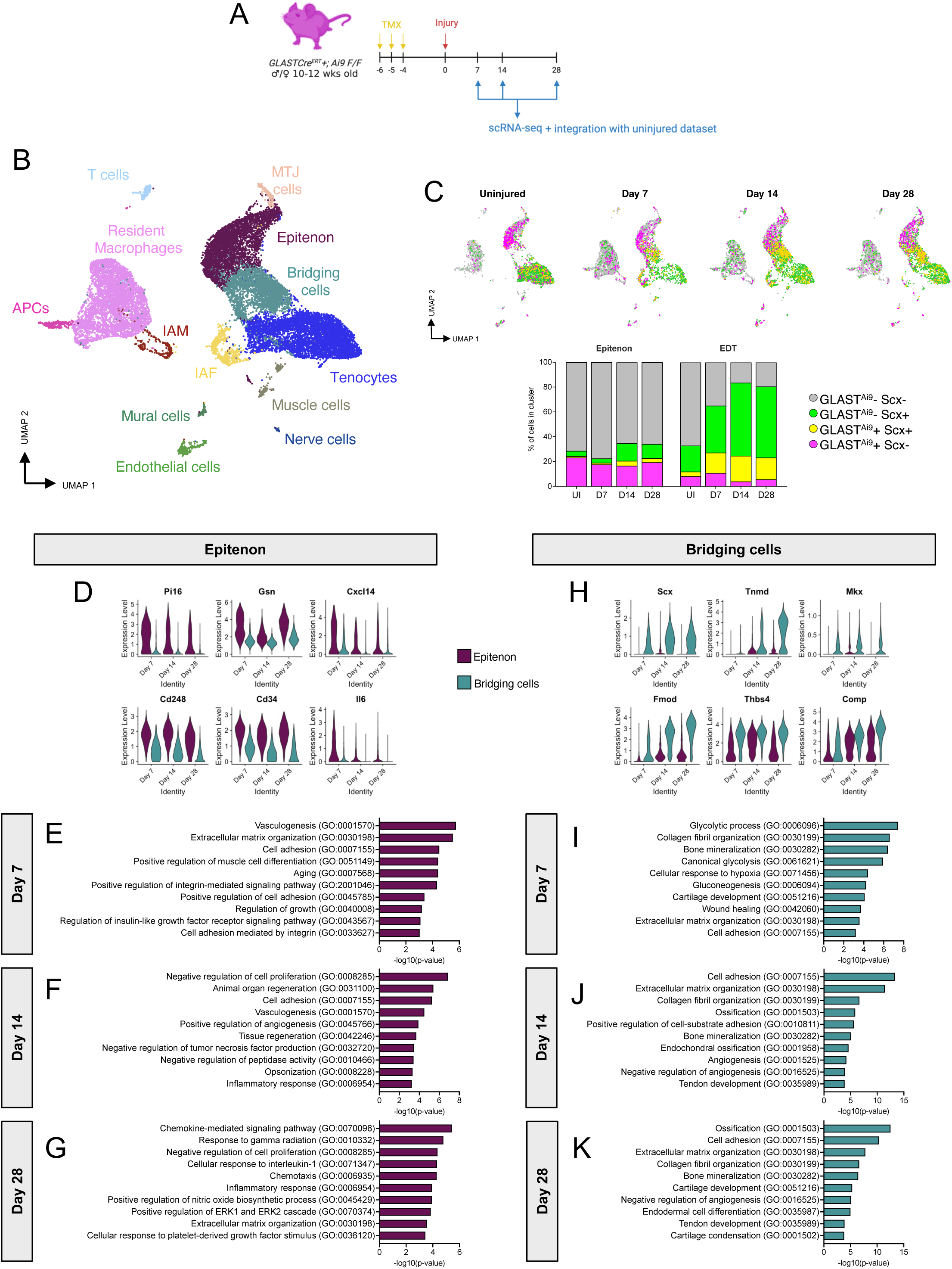
Epitenon-derived cells exhibit dual fates during healing. (**A**) Overview of experimental design. (**B**) Annotated UMAP plot of integrated healing dataset containing cells from uninjured FDL tendon and tendons at 7, 14, and 28 days post-injury. IAF = Injury-associated fibroblasts, APCs = Antigen-presenting cells. IAM = Injury-associated macrophages. (**C**) UMAP plots (top) and quantification (bottom) of epitenon cells (magenta: GLAST^Ai9^+; Scx-), tenocytes (green: GLAST^Ai9^-; Scx+), and Bridging cells; yellow: GLAST^Ai9^+; Scx+) in the integrated dataset at each timepoint. (**D** - **E**) Increased expression of fibrotic markers in the epitenon cluster (**D**) and tenogenic markers in the Bridging cells cluster (**E**) over time. (**F** - **K**) GO terms enriched in the epitenon (**F - H**) and Bridging cells (**I** - **K**) clusters at each timepoint.

To characterize the overall contribution of the epitenon and bridging cell clusters to tendon healing, we examined DEG in each cluster over time (**Fig. S4E-J**). This revealed a generalized healing response in both clusters, with enrichment for biological processes broadly associated with collagen deposition and ECM organization at each timepoint post-injury. Of note, a distinct pro-inflammatory response represented by enrichment for pathways associated with interleukin (IL)-1/ IL-1 beta was observed in the epitenon cell cluster at D7 (**Fig. S4D, S4E**) but was completely absent in the bridging cell cluster (**Fig. S4K**), suggesting that epitenon-derived cells play a role in initiating the early inflammatory response. To determine cell-type specific functions against this background of general healing, we performed DEG analysis to directly compare the epitenon and bridging cell clusters at each timepoint (**Supplemental Table 2**).

Consistent with the idea of dual fates for epitenon-derived cells, the epitenon cluster displayed significantly higher expression of pro-inflammatory/profibrotic genes, including C-X-C motif chemokine ligand 14 (*Cxcl14*), peptidase inhibitor 16 (*Pi16*), gelsolin (*Gsn*)*, Cd248, Cd34,* and *Il6* (**Fig. 4D**), at all timepoints compared to the bridging cell cluster. Gene ontology (GO) analysis of DEG in the epitenon cluster at D7 post-injury showed enrichment for pathways associated with vasculogenesis, ECM organization, cell adhesion/integrin-mediated signaling, and the insulin-like growth factor signaling pathway (a known regulator of collagen synthesis in tendon/ligament healing)^26–28^ (**Fig. 4E**), supporting a role for epitenon cells as early orchestrators of the healing response beyond an initiation of the inflammatory phase. At D14, epitenon cells were enriched for processes that suggested a shift towards a pro-fibrotic/pro-inflammatory phenotype, including positive regulation of angiogenesis/vasculogenesis, cell adhesion, and inflammatory responses (**Fig. 4F**). At D28, epitenon cells were largely pro-inflammatory, with enrichment for processes including chemokine-mediated signaling, cellular response to IL-1, chemotaxis, inflammatory response, ECM organization, and cellular response to PDGF stimulus (**Fig. 4G**). Given that fibrotic scar formation is thought to be due to the prolonged activity of activated fibroblast populations, the pro-inflammatory phenotype of epitenon cells at D28 (well into the remodeling phase of healing) and enrichment for pathways associated with fibrovascular processes (angiogenesis/vasculogenesis, chemokine-mediated signaling) combined with localization at the periphery of the injury site suggests that epitenon-derived cells are involved in the formation of peritendinous scar tissue. In contrast to the epitenon cluster, the bridging cell cluster displayed significantly higher levels of the tenogenic genes *Scx*, tenomodulin (*Tnmd*), Mohawk (*Mkx*), fibromodulin (*Fmod*), thrombospondin-4 (*Thbs4*), and cartilage oligomeric matrix protein (*Comp*) at each timepoint post-injury (**Fig. 4H**). At D7, GO terms were largely associated with glycolysis (glycolytic processes, canonical glycolysis, gluconeogenesis), ECM/collagen fibril organization, and wound healing. Enrichment for pathways associated with musculoskeletal cell differentiation (bone mineralization and cartilage development) similarly point to the bridging cell as a priming progenitor cell population at D7. At both D14 (**Fig. 4J**) and D28 (**Fig. 4K**), the bridging cell cluster was enriched for processes indicative of ECM/collagen fibril organization, musculoskeletal cell differentiation (ossification, endochondral ossification, tendon development, cartilage development), and negative regulation of angiogenesis, suggestive of a pro-regenerative, tenogenic injury response.

To confirm these differing fate trajectories suggested by our *in silico* analysis, we performed immunofluorescent staining for the pro-inflammatory/pro-fibrotic cytokine CXCL14^29, 30^ (**Fig. 5A**), which was highly expressed throughout healing in the epitenon cell cluster compared to the bridging scar cluster (**Fig. 4D**). Consistent with its role as a pro-inflammatory cytokine, the number of CXCL14+ cells in the bridging scar tissue peaked at D7 during the inflammatory phase before declining through D28 post-injury (**Fig. 5B**: D3: 16.8 ± 5.9%, D7: 78.4 ± 16.3%, D10: 70.9 ± 14.6%, D14: 37.9 ± 9.5%, D21: 19.0 ± 4.7%, D28: 30.0 ± 7.3%). In contrast, the number of CXCL14+ cells in the peripheral capsule surrounding the injury site remained consistently high throughout the timecourse examined (**Fig. 5C**: D3: 38.2 ± 8.6%, D7: 29.7 ± 18.8%, D10: 45.3 ± 12.1%, D14: 55.1 ± 4.1%, D21: 44.7 ± 7.0%, D28: 54.8 ± 17.8%). In addition, the percent of GLAST^Ai9;^ CXCL14+ in the peripheral capsule was significantly higher at every timepoint compared to those in the bridging capsule (D3: p = 0.0023, D7: p= 0.0282, D10: p = 0.0104, D14: p = 0.0061, D21: p = 0.0008, D28: p = 0.0283). Combined with increased *de novo* expression of Scx^GFP^ by epitenon-derived tenocytes in the bridging tissue (**Fig. 3**), these data support spatially-distinct fates for epitenon cells during tendon healing.

**Fig 5.**
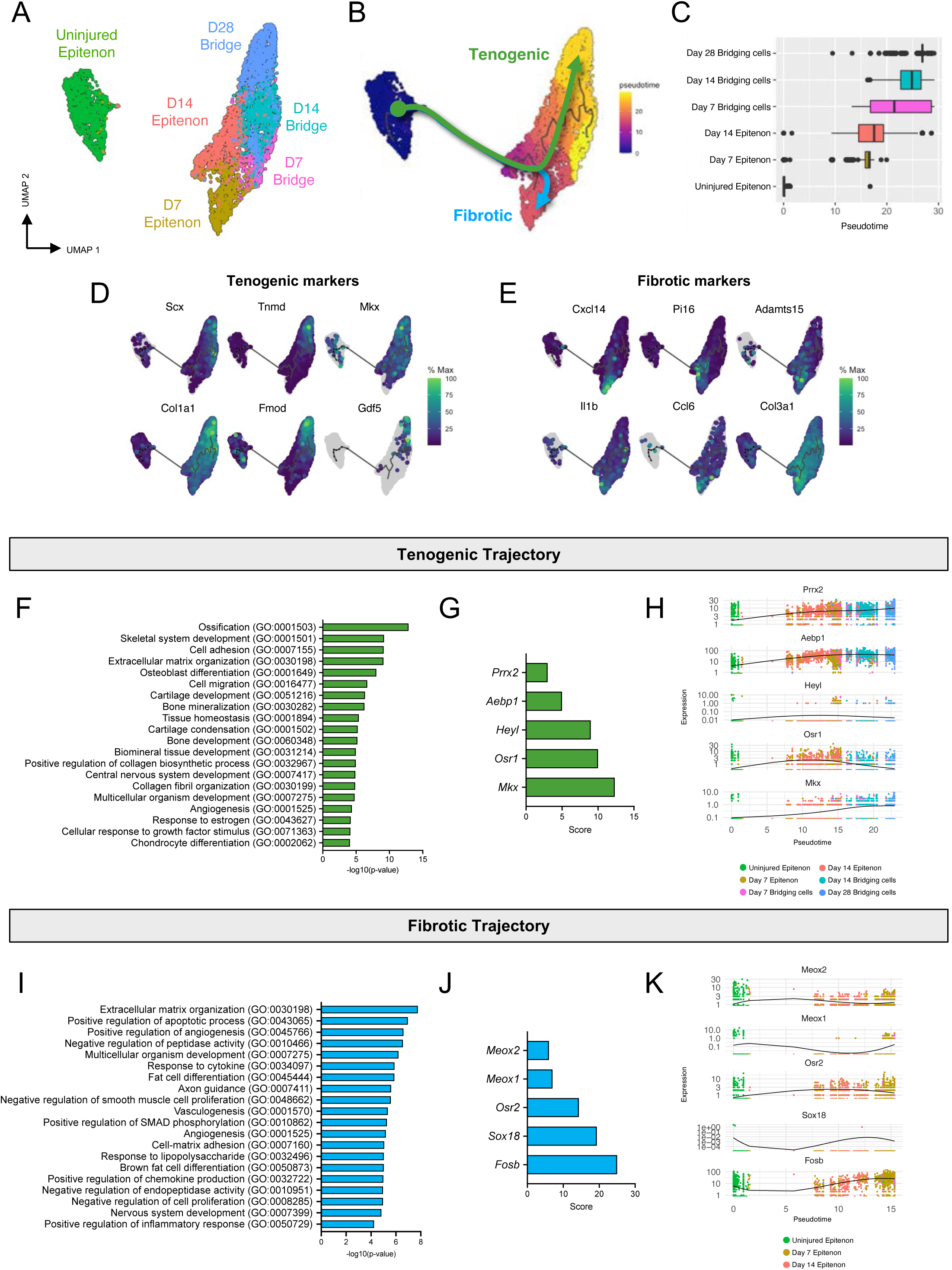
GLAST^Ai9^ epitenon-derived cells on the periphery of the injury site exhibit increased expression of the pro-inflammatory/pro-fibrotic cytokine CXCL14. (**A**) Representative images of healing tendons from GLAST^Ai9^; Scx^GFP^ mice at days 3, 7, 10, 14, 21, and 28 post-injury showing the presence of GLAST^Ai9^ cells (red) and CXCL14 (yellow). Tendon stubs are outlined by white dashed lines and the scar area is outlined by white dotted lines. Scale bars = 200 μm, magnified image scale bars = 50 μm. Pink outlined magnified images denote GLAST^Ai9^ cells on the periphery of the injury site and light blue outlined magnified images show GLAST^Ai9^ cells within the bridging scar tissue. Quantification of GLAST^Ai9^; CXCL14+ cells in the bridging scar tissue (**B**) vs the peripheral capsule (**C**). Bars represent the mean cell number and error bars denote the standard deviation. In all images, nuclei are counterstained with Hoechst (blue). All images are representative of n = 3-4 mice per timepoint.

### Pseudotime analysis reveals both tenogenic and fibrotic trajectories of epitenon cells

In addition to the inclusion of multiple timepoints in our scRNA-seq analysis, we utilized pseudotime trajectory analysis to better understand the progression of epitenon-derived cells to either a fibrotic or tenogenic fate post-injury. For this analysis, we subset the uninjured epitenon (origin) cluster, the D7, D14, and D28 bridging cell clusters, and the D7 and D14 epitenon clusters (**Fig. 6A**). The D28 epitenon cluster was removed as it appeared earliest in pseudotime compared to the other post-injury epitenon clusters (**Fig. S6A, S6B**), indicating that in addition to being distinctly pro-fibrotic (**Fig. 4**, **5**), a sizable proportion of epitenon cells at D28 were returning to a more homeostatic state consistent with the natural resolution/reduction of peritendinous scar tissue in this model at this time. In support of a fate bifurcation at ∼D14, pseudotime trajectory analysis inferred a branched trajectory (**Fig. 6B**) from the uninjured epitenon through the injured epitenon (**Fig. 6C**) to either the D28 bridging cell (tenogenic) or the D7/D14 epitenon (fibrotic). Correspondingly, tenogenic markers including *Scx*, *Tnmd*, *Mkx*, *Col1a1*, *Fmod*, and growth differentiation factor 5 (*Gdf5*) were significantly upregulated along the tenogenic trajectory (**Fig. 6D**) whereas fibrotic markers including *Cxcl14*, *Pi16*, ADAM metallopeptidase with thrombospondin type 1 motif 15 (*Adamts15*), *Il1b*, chemokine (C-C motif) ligand 6 (*Ccl6*), and collagen type 3 alpha 1 (*Col3a1*) were significantly upregulated along the fibrotic trajectory in pseudotime (**Fig. 6E**). Similar upregulation of tenogenic (**Fig. S6C**) and fibrotic (**Fig. S6D**) marker genes in real-time support the pseudotime lineage reconstruction.

**Fig. 6.**
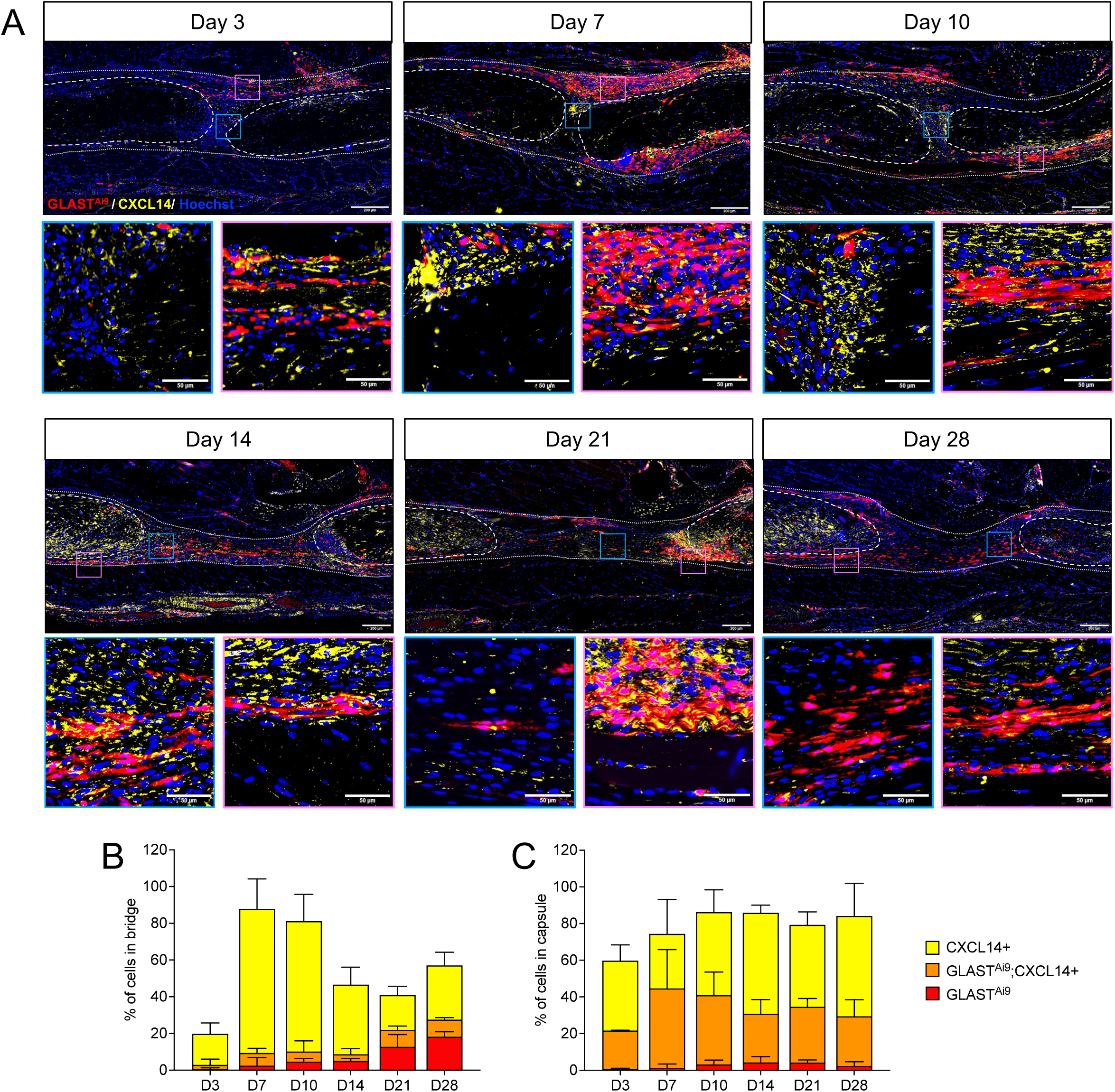
Branched trajectory analysis and transcriptional regulation of epitenon-derived cell fates. (**A**) Annotated UMAP graph of clusters used for trajectory analysis. (**B**) UMAP depicting inferred branched trajectory from uninjured epitenon to tenogenic (green) and fibrotic (blue) fates. (**C**) Pseudotime progression of epitenon-derived cells from uninjured epitenon to day 28 EDT. (**D** - **E**) Upregulation of tenogenic (**D**) and fibrotic (**E**) markers along the different fate trajectories. (**F**) GO terms enriched along in the tenogenic trajectory. (**G**) Putative transcription factors regulating tenogenesis of epitenon cells. (**H**) Potential tenogenic transcriptional regulators in pseudotime, colored by cell type in real time. (**I**) GO terms enriched along in the fibrotic trajectory. (**J**) Putative transcription factors regulating fibrotic fate of epitenon cells. (**K**) Potential fibrotic transcriptional regulators in pseudotime, colored by cell type in real time.

To gain insight into the factors regulating the tenogenic and fibrotic fates of epitenon cells, we examined gene modules (groups of co-regulated genes) for each trajectory branch. Consistent with tenogenic differentiation from a progenitor cell population, GO term analysis of the tenogenic gene module revealed enrichment for processes associated with musculoskeletal development (*e.g.,* ossification, skeletal system development, osteoblast and chondrocyte differentiation, bone and cartilage development), collagenous ECM deposition (*e.g.,* ECM organization, positive regulation of collagen biosynthetic process, collagen fibril organization), and cellular response to growth factor stimulus (**Fig. 6F, Supplemental Table 3**). Transcription Factor Enrichment Analysis (TFEA) of the tenogenic gene module identified paired related homeobox 2 (*Prrx2*), AE binding protein 1 (*Aebp1*), hes-related family bHLH transcription factor with YRPW motif-like (*Heyl*), odd-skipped related transcription factor 1 (*Osr1*), and *Mkx* as likely transcriptional regulators of the tenogenic fate (**Fig. 6G**, **6H**).

GO term analysis of the fibrotic gene module showed enrichment for biological processes associated with ECM organization/cell-matrix adhesion, pro-angiogenic/neurogenic processes (*e.g.,* positive regulation of angiogenesis, axon guidance, vasculogenesis, nervous system development), and pro-inflammatory processes (*e.g*., response to cytokine, response to lipopolysaccharide, positive regulation of chemokine production, positive regulation of inflammatory processes), all processes supporting involvement of the epitenon in the formation of fibrovascular scar (**Fig. 6I, Supplemental Table 3**). TFEA identified mesenchyme homeobox (*Meox*) 2*, Meox1*, *Osr2*, SRY-box transcription factor 18 (*Sox18*), and FBJ murine osteosarcoma viral oncogene homolog B (*Fosb*) as potential regulators of the pro-fibrotic fate (**Fig. 6J**, **6K**).

### Ablation of epitenon-derived cells post-injury improves functional recovery

Given the pro-fibrotic phenotype of epitenon-derived cells on the periphery of the healing site which implicates them in the formation of function-limiting peritendinous scar, we next sought to determine whether ablation of GLAST^Ai9^ epitenon-derived cells following tendon injury would improve functional recovery. To ablate epitenon-derived cells, we generated GLAST^Ai9;DTR^ mice in which diphtheria toxin receptor (DTR)-expressing GLAST^Ai9^ epitenon cells (labeled prior to injury; **Fig. 7A**) undergo apoptosis after exposure to diphtheria toxin (DT).This model resulted in >80% depletion of GLAST^Ai9^ epitenon cells in uninjured tendons that persisted to at least day 5 (>70%) post-depletion (**Fig. S7**). GLAST^Ai9;DTR^ mice underwent tendon injury and repair surgery and on days 7-10 post-injury (the peak of GLAST^Ai9^ cell proliferation), mice received local DT or control (PBS) injections directly to the injury site. Tendons were then assessed for functional (metatarsophalangeal joint range of motion [MTP ROM] and gliding) and biomechanical (maximum load and stiffness) assessment at days 14 and 28 post-injury (**Fig. 7A**). At both 14 and 28 days post-injury (**Fig. 7B**), DT treated tendons exhibited significantly increased MTP ROM compared to PBS treated tendons (**Fig. 7C**; D14-PBS: 29.85 ± 13.29°, DT: 43.01 ± 15.72°, p = 0.0316; D28-PBS: 46.19 ± 8.27°, DT: 60.36 ± 7.72°, p = 0.0026) and significantly decreased gliding resistance (**Fig. 7D**; D14-PBS: 38.58 ± 21.27, DT-19.46 ± 13.84, p = 0.0113; D28-PBS: 15.20 ± 5.91, DT: 8.23 ± 1.63, p = 0.0113), indicative of improved functional recovery due to decreased peritendinous scar. No significant differences were observed in the maximum load (**Fig. 7E**; D14-PBS: 0.66 ± 0.25 N, DT: 0.85 ± 0.30 N, p = 0.1889; D28-PBS: 2.04 ± 0.86 N, DT: 2.10 ± 0.61 N, p = 0.8712) or stiffness (**Fig. 7F**: D14-PBS: 1.6 ± 1.06 N/mm, DT: 2.40 ± 0.86 N/mm, p = 0.0588; D28-PBS: 2.93 ± 1.48 N/mm, DT: 4.17 ± 3.42 N/mm, p = 0.5490) of GLAST^Ai9^ cell ablated tendons compared to controls. Combined, these data implicate GLAST^Ai9^ epitenon cells in the formation of peritendinous scar and suggest that manipulation of their presence post-injury can significantly improve functional recovery without disrupting the tendon healing process.

**Fig. 7:**
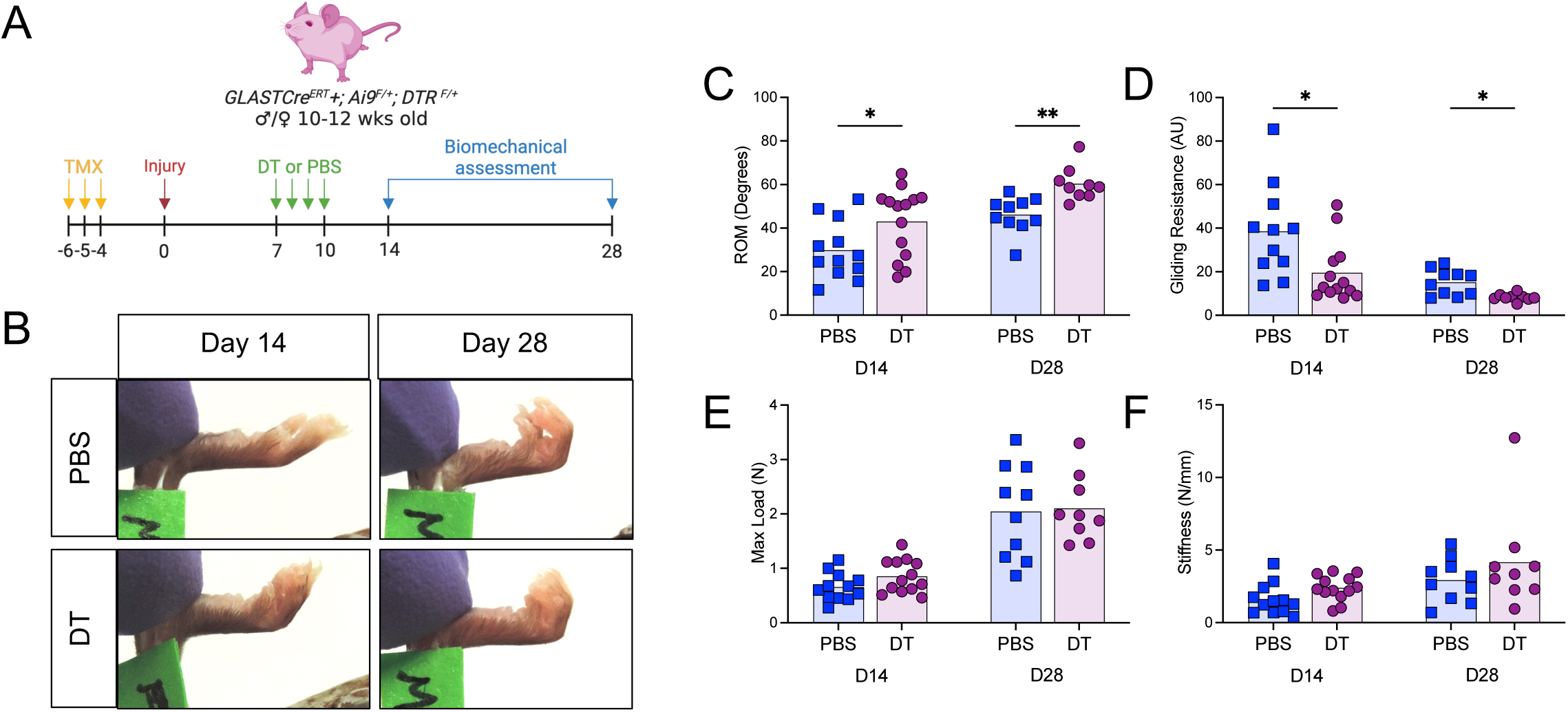
Depletion of GLAST^Ai9^ epitenon cells improves functional recovery. (**A**) Overview of experimental design. (**B**) Representative images of metatarsophalangeal joint (MTP) range of motion (ROM) from PBS (control) and diphtheria toxin (DT) treated tendons at days 14 and 28 post-injury. Quantification of the mean (**C**) MTP ROM, (**D**) gliding resistance, (**E**) maximum load at failure, and (F) stiffness of PBS or DT treated tendon repairs at days 14 and 28 post-injury. n= 10-14 mice per timepoint per group. *p<0.05 **p<0.01

### Human peritendinous adhesion cells are transcriptomically similar to pro-fibrotic mouse epitenon cells

In order to determine the relationship between epitenon-derived pro-fibrotic cells in our mouse model and the cell populations present in mature human peritendinous scar, we integrated our mouse healing dataset with scRNA-seq datasets derived from human flexor tendon peritendinous scar tissue removed during tenolysis surgery, both from our lab and from a previous publication^30^ (**Fig. 8A**). A total of 42,596 human peritendinous scar cells were used for downstream analyses. First, cell populations in the integrated human dataset were annotated based on established markers (**Fig. S8A**), with all cell populations being present in each sample in varying proportions (**Fig. S8B**). The mouse healing integrated UMAP was then used as a reference and human cells were mapped to their corresponding mouse clusters (**Fig. 8B**). Consistent with our hypothesis that the epitenon contributes cells to the peritendinous scar tissue, the majority of human scar cells mapped to the mouse epitenon cell cluster (28.1%; **Fig. 8B**). Finally, to determine whether human peritendinous scar cells may be derived from epitenon cells, we interrogated the mouse scRNAseq-dataset for epitenon-specific markers that were conserved throughout healing. *Ccn3* is not only expressed specifically in the mouse epitenon during homeostasis, it is also expressed by epitenon cells throughout healing (**Fig. 8C**, **8D**) and by fibroblasts present in human peritendinous scar tissue (**Fig. 8E**, **8F, S8C**). Confirmatory immunofluorescent staining for CCN3 in uninjured human flexor tendon (**Fig 8G**) and scar tissue from an additional three tenolysis patients revealed the presence of CCN3+ cells throughout. Combined, these data suggest that the fibroblasts present in established human peritendinous scar tissue may originate from the epitenon.

**Fig. 8.**
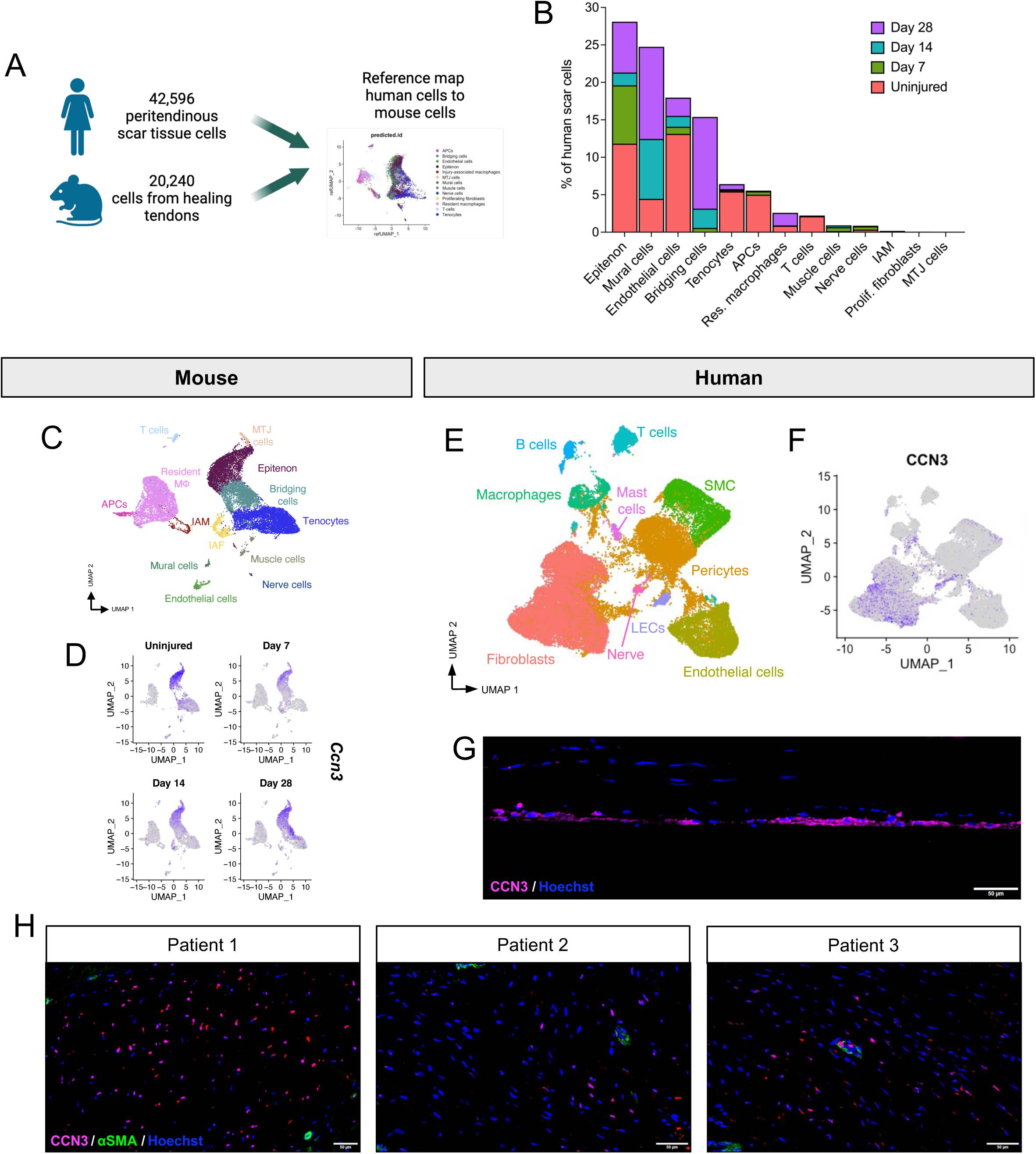
Integration of mouse healing scRNA-seq dataset with human peritendinous scar tissue. (**A**) Experimental overview. (**B**) Proportion of human peritendinous scar cells that correspond to individual mouse clusters, colored by timepoint. IAF = Injury-associated fibroblasts, APCs = Antigen-presenting cells. IAM = Injury-associated macrophages. (**C**) UMAP of annotated integrated mouse healing scRNA-seq dataset. (**D**) Feature plots showing conserved expression of Ccn3 in the mouse epitenon throughout healing. (**E**) UMAP of annotated integrated human peritendinous scar scRNA-seq dataset. (**F**) Feature plot showing expression of CCN3 in the fibroblast cluster. (**G**) Immunofluorescent staining of CCN3 (magenta) in uninjured human flexor tendon. Scale bar = 50 μm. (**H**) Immunofluorescent staining for conserved epitenon-derived cell marker CCN3 (magenta) in the peritendinous scar tissue from three individual human patients. In all immunofluorescent images, nuclei are counterstained with Hoechst (blue).

## DISCUSSION

Understanding the specific identity, origin, and fate of the various cell populations that contribute to tendon healing is vital to the development of targeted therapeutics to improve the naturally poor healing response of tendons. Here, we report an in-depth spatial and temporal characterization of the epitenon-derived cellular contribution to tendon healing and identify the epitenon as a niche for a bi-fated progenitor cell population that participates in both tendon regeneration and peritendinous fibrosis following acute flexor tendon injury. Depletion of pro-fibrotic epitenon cells post-injury significantly improved functional recovery, and integrated scRNA-seq analysis of mouse tendon healing with human peritendinous scar tissue found striking transcriptional similarity between pro-fibrotic mouse epitenon cells and the fibroblasts present in human scar tissue, implicating the epitenon as a major contributor to the formation of peritendinous fibrosis. Combined, these data establish the epitenon as the cellular origin of an important progenitor cell population that could be leveraged to improve overall tendon healing via modulation of both peritendinous scar formation and regeneration of the tendon itself (**Fig. 9**).

**Fig 9.**
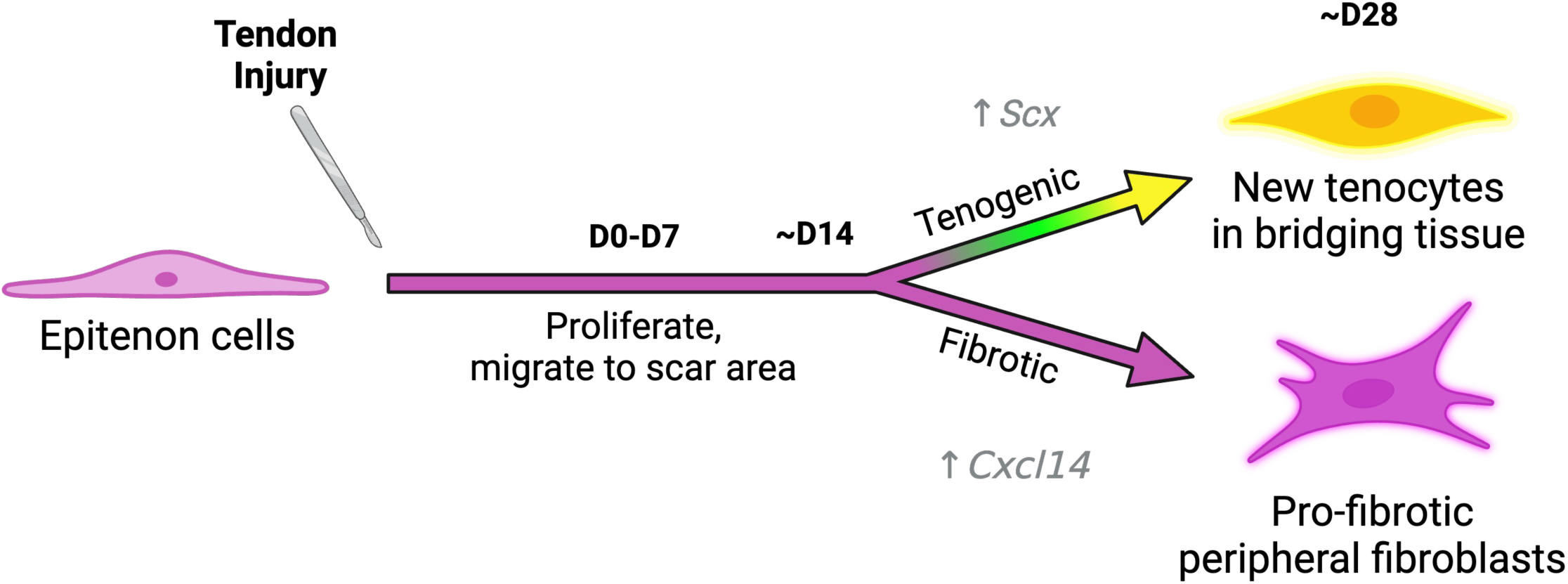
Working model of the epitenon cell contribution to flexor tendon healing.

The initial goal of this study was to identify a specific genetic driver that could be used to determine the epitenon cell contribution to the formation of peritendinous adhesions following acute flexor tendon injury. To that end, we have established the *GLASTCre^ERT^*driver as a tool to target the epitenon in adult mice and identified both GLAST and CCN3 as conserved epitenon-specific markers in mice and humans during both homeostasis and throughout healing. This finding will greatly enhance future work to understand the overall cellular landscape of tendon healing across models. CCN3 (also known as nephroblastoma overexpressed or *Nov*) is a member of the CCN family of extracellular matrix-associated signaling proteins that has demonstrated involvement in many cellular processes, including angiogenesis^31^, bone regeneration^32–34^, myogenesis^35, 36^ and fibrosis where it has shown to have both pro-^37^ and anti-fibrotic effects^38–40^. Aside from a single report of high CCN3 expression in developing tendons^41^ we are unaware of any studies examining the role of CCN3 in tendon. Given the crucial role that CCN3 and other CCN family members play in myogenesis and bone regeneration, additional work is warranted to understand potential contributions of CCN3 to tendon homeostasis and healing.

In addition, GLAST^Ai9^ epitenon cells localize to the peripheral scar tissue during murine tendon healing and have a distinctly pro-inflammatory/pro-fibrotic transcriptomic profile that was highly similar to that of fibroblasts isolated from human peritendinous scar tissue. Consistently, depletion of GLAST^Ai9^ epitenon cell post-injury significantly improved functional recovery without impacting biomechanical properties of the healing tendon. Combined, these results suggest that the epitenon plays an integral role in the development of peritendinous adhesions. Surprisingly, in addition to being a source of pro-fibrotic cells during healing, GLAST^Ai9^ epitenon-derived cells also underwent tenogenesis in the organized bridging tissue post-injury, indicating that the epitenon serves as a niche for a bi-fated progenitor cell population. Previous studies have identified a number of potential adult tenogenic progenitor cell populations derived from various parts of the tendon that participate in tendon healing, including those derived from the tendon sheath/paratenon (patellar tendon: α*SMA+*^42^, *Tppp3+*^43^; tibialis anterior tendon: *Bglap+*^44^) and parenchyma (Achilles tendon: *Axin2+*^45, 46^, *Scx+*^47^; FDL tendon: *Scx*+^20^). The degree to which these populations are specific to different compartments appears to vary across tendons: for example, *Tppp3* is a sheath/paratenon-specific marker in the patellar tendon but not in the Achilles^45, 48^ and we observed very little GLAST^Ai9^ labeling of the patellar tendon epitenon. This suggests that different progenitor populations contribute to tendon healing in anatomically distinct tendons. Additional work is required to determine which progenitor populations are present and contribute to tendon healing in different tendons and how these populations might be leveraged to improve healing in a tendon-specific or injury-specific manner.

An important aspect of tendon healing highlighted by this study is the degree to which both temporal and spatial cell fates govern the overall healing response. Our current and previous work using this model has demonstrated that there are several fate trajectories that occur post injury: 1) tenocytes that remain within the tendon stubs undergo activation to a transient myofibroblast phenotype that helps to remodel the existing tendon tissue to integrate with the newly formed bridging scar tissue^20, 23^, 2) intrinsically and extrinsically-derived cells migrate to the injury site and form the bridging scar tissue between the cut ends of the tendon, some of which become new tenocytes post-injury^20, 49, 50^, and 3) cells from the epitenon contribute to the formation of fibrotic peripheral capsule (this study). The degree to which these processes influence each other is unknown but is an interesting question for future studies as broadly targeting “fibrosis” is likely to be insufficient or lead to undesirable outcomes.

In addition to the epitenon, a small number of GLAST^Ai9^ cells were found within the body of the tendon during homeostasis (∼9% of all tenocytes). As other studies have described progenitor cell populations within the tendon body that contribute to tendon healing^20, 45–47^, it is possible that the GLAST^Ai9^ tenocytes also represent a tendon body-derived progenitor cell population; however, this does not appear to be the case. A similar proportion (11-13%) of GLAST^Ai9^ cells continue to cluster with the tenocyte cluster prior to injury and throughout healing, which suggests that they share a similar post-injury fate to tenocytes and remain transcriptomically distinct from GLAST^Ai9^ cells found in either the bridging cell or epitenon cell clusters. Moreover, we consistently observed a number of GLAST^Ai9^; IllSMA+ myofibroblasts located within the tendon stubs (Supplemental Fig. 3A), indicating that GLAST^Ai9^ tenocytes contribute to the tenocyte-derived myofibroblast population post-injury. Additional work is required in order to understand whether there is a relationship between GLAST^Ai9^ epitenon cells and GLAST^Ai9^ tenocytes during tendon homeostasis, or whether GLAST^Ai9^ tenocytes may represent a specialized subpopulation of tenocytes.

Interestingly, while we observed that approximately 30% of ScxGFP+ tenocytes in the bridging scar tissue arise from GLAST^Ai9^ epitenon cells, depletion of this population (as part of the broader GLAST^Ai9^ cell depletion post-injury) did not significantly affect bulk mechanical properties. Given that the remaining 70% of ScxGFP+ tenocytes in the bridge arise from other (non-GLAST-lineage) sources, this suggests that GLAST^Ai9^ epitenon cells are not the predominant tenogenic progenitor population that responds to acute injury. This could explain why loss of this tenogenic progenitor does not appear to impact bulk mechanical properties; however, the relationship between the presence/absence of tenogenic progenitors and mechanical properties is not clear. For example, previous work from our group demonstrated that broad depletion of tendon cells using the Scx-Cre driver unexpectedly improved bulk mechanical properties^59^. Similarly, we observed a slight increase in stiffness at D14 with GLAST cell depletion (though not statistically significant: p = 0.0588) in this study. Combined, these findings suggest that it is not just the presence or absence of particular cell populations, but also altered stimulatory/inhibitory signaling between populations that ultimately dictates healing outcomes. Additional work is required to understand the complex interplay between cell populations to identify strategies that effectively promote regenerative healing.

This study demonstrates the power of combined lineage tracing and scRNA-seq to gain unprecedented insight into both cellular behavior and origins. In addition, the combined use of animal models with patient-derived samples highlights the enormous potential to identity novel markers and translational mechanisms that would not be possible using human tissue alone. For example, despite the recent proliferation of scRNA-seq studies using human tendon^51–54^, annotation of distinct cell clusters has remained difficult due to a lack of specific markers for various tendon cell populations. In particular, lack of an epitenon cell marker has meant that this important structure has been entirely excluded from human transcriptomic studies. Here, through integrated dual species scRNA-seq analysis, we uncovered epitenon-specific markers that can be used to annotate the epitenon in future investigations. Additionally, through analysis of conserved markers for pro-fibrotic epitenon-derived cells, we identified epitenon-derived cells as a novel cellular therapeutic target to reduce peritendinous adhesion formation in humans following acute flexor tendon injures.

In summary, these data identify the epitenon as a source of bi-fated progenitor cells that contribute to both tendon fibrosis and regeneration in mice. Moreover, we provide evidence that this role is partially conserved in human tendon healing, as epitenon-specific markers were present in a small cohort of patient-derived peritendinous scar tissue. Identifying the specific role and fates of epitenon cells during tendon healing is a significant step towards the ultimate goal of improved tendon healing through the suppression of adhesion formation and promotion of tendon regeneration.

### Limitations of the study

The main limitation of this relates to the use of the *GLASTCre^ERT^*driver for lineage tracing experiments. Both *in situ* imaging and scRNA-seq in intact mouse flexor tendons showed that not all epitenon cells are targeted by the *GLASTCre^ERT^*driver and therefore only a subset of epitenon cells (∼24%) are able to be identified spatially or manipulated during tendon healing. Additionally, the *GLASTCre^ERT^*driver targets several non-epitenon cell populations, including adipocytes, perivascular cells, and macrophages, therefore we cannot rule out the contribution of extrinsically-derived GLAST^Ai9^ cells to the tendon healing process by lineage tracing alone. To overcome this, we combined genetic lineage tracing with scRNA-seq which facilitated identification of both epitenon- and epitenon sub-population-specific markers. Future investigations utilizing novel Cre drivers generated using these markers will further clarify the specific functional roles of all epitenon cell populations to tendon healing. An additional limitation of the current study design is the exclusion of timepoints between uninjured tendon and day 7 post-surgery for scRNA-seq. Due to this, the initial factors or cellular signals that activate epitenon cells post-injury remain to be determined. Future studies incorporating these early timepoints will be critical in determining the temporal dynamics and molecular signals that govern the early stages of epitenon cell activation and their subsequent differentiation pathways.

In addition, there is much confusion in the literature regarding the naming of different parts of the tendon (sheath vs epitenon vs paratenon), due largely to a lack of markers for each compartment and the presumption that all tendons are compositionally the same. Discrepancies in how different tendon compartments are targeted with various drivers in anatomically distinct tendons suggests several important things: 1) tendons are not structurally identical^55, 56^, 2) cell populations present in tendon compartments (e.g. epitenon, paratenon, sheath, etc.) vary between different tendons, and 3) progenitors involved in tendon healing differ between tendons (i.e. flexor vs patellar). As such, it is crucially important for the field to identify and characterize different tendon cell populations in anatomically distinct tendons. This study is a first step towards that goal as we can now begin to answer some of the important outstanding questions in the field (e.g. Do progenitor cell populations differ between tendons? To what degree are specific cellular populations and/or fibrotic processes conserved between tendons?) While answering these questions is outside the scope of the present study, the tools and markers identified here will help enable these future investigations.

## METHODS

### Resource Availability

#### Contact for Reagent and Resource Sharing

Further information and requests of reagents can be directed to and fulfilled by the Lead Contact, Anne Nichols.

### Materials availability

This study did not generate new unique reagents.

### Data availability

Single-cell RNA-sequencing data have been deposited at GEO and are publicly available as of the date of publication (Accession number TBD prior to publication). Microscopy data reported in this paper will be shared by the lead contact upon request. Any additional information required to reanalyze the data reported in this paper is available from the lead contact upon request.

### Code availability

This paper does not report original code.

### Experimental model and subject details

#### Animal Ethics

All experiments were carried out in strict accordance with the recommendations in the Guide for the Care and Use of Laboratory Animals of the National Institutes of Health. All animal procedures were approved by the University Committee on Animal Research (UCAR) at the University of Rochester (protocol #2017-004E). Mice were kept in standard housing conditions throughout breeding and experimentation.

#### Mice

*GLASTCre^ERT^* (STOCK Tg[Slc1a3-cre/ERT]1Nat/J; #012586)^12^, *ROSA-Ai9* (B6.Cg-Gt[ROSA]26Sortm9[CAG-tdTomato]Hze/J; #007909)^16^, and ROSA-iDTR (C57BL/6-Gt(ROSA)26Sortm1(HBEGF)Awai/J; # 007900) mice were obtained from The Jackson Laboratory (Bar Harbor, ME, USA). *Scleraxis (Scx)-GFP* (Tg[Scx-GFP]1Stzr/Tg[Scx-GFP]1Stzr; referred to as Scx^GFP^)^19^ reporter mice were generously provided by Dr. Ronen Schweitzer (Oregon Health Sciences University). To identify and trace GLAST-lineage cells by immunofluorescence and scRNA-seq, *GLASTCre^ERT^* mice were crossed to the *ROSA-Ai9* strain to generate experimental animals (*GLASTCre^ERT+^; Ai9^F/F^*: referred to as GLAST^Ai9^). Male and female mice GLAST^Ai9^ mice (10-12 weeks old) received three i.p. injections of tamoxifen (TMX; 100 mg/kg in corn oil, T5648 and C8267, Sigma Aldrich, St. Louis, MO, USA) and tendons were harvested 14 days after the final injection. To evaluate potential tenogenic differentiation of GLAST-lineage cells, GLAST^Ai9^ mice were crossed to Scx^GFP^ mice (*GLASTCre^ERT^; Ai9^F/+^; Scx^GFP+^*: referred to as GLAST^Ai9^; Scx^GFP^) which allows tracking of cells that actively express the tenogenic transcription factor scleraxis (*Scx*). GLAST^Ai9^; Scx^GFP^ mice received the same TMX regimen described above for GLAST^Ai9^ mice. To facilitate depletion of GLAST-lineage cells post-injury, GLAST^Ai9^ mice were crossed to ROSA-iDTR^F/F^ (referred to as DTR) to generate experimental animals (*GLASTCre^ERT+^; Ai9^F/+^; DTR^F/+^*: referred to as GLAST^Ai9;DTR^).

#### Flexor tendon injury model

To trace GLAST^Ai9^ epitenon cells (GLAST^Ai9^, GLAST^Ai9^; Scx^GFP^, and GLAST^Ai9;DTR^) and induce expression of the diphtheria toxin receptor (DTR) in GLAST^Ai9^ cells (GLAST^Ai9;DTR^), male and female mice (10-12 weeks old) received three i.p. TMX injections beginning seven days prior to flexor tendon injury, allowing a four-day TMX washout period. This labeling regimen ensured that only cells expressing GLAST prior to injury were labeled and that no subsequent labeling occurred during tendon healing. Mice then underwent complete transection and repair of the flexor digitorum longus (FDL) tendon in the right hind paw as previously described.^22, 23, 57–61^. Briefly, mice were anesthetized with Ketamine (100mg/kg) and Xylazine (10mg/kg) and sustained-release buprenorphine (ZooPharm™; 0.5-1.0 mg/kg) was applied subcutaneously for post-operative pain management. To reduce chances of rupture at the repair site, the FDL tendon was first transected at the myotendinous junction (MTJ) and the skin was closed with non-resorbable 5-0 suture (#668G Ethilon, Ethicon Inc., Bridgewater, NJ, USA). Transection of the MTJ results in a transient decrease in loading, with reintegration of the MTJ observed by day 7-10 post-surgery in this model. Following sterilization of the surgery site, a small incision was made on the plantar surface of the right hind paw. The FDL tendon was carefully located, completely transected at the mid substance, and repaired using non-resorbable 8-0 suture (#2808G Ethilon, Ethicon Inc.) using a modified Kessler pattern. Skin over the surgery site was closed with 5-0 suture. Mice were carefully monitored for 72 hours post-surgery to ensure their return to prior cage activity, food intake, and water consumption.

#### GLAST^Ai9^ cell depletion

On days 7 through 10 post-injury, GLAST^Ai9; DTR^ mice (n = 9-14 per group per timepoint) received localized injections of diphtheria toxin (DT) into the right hind paw (20 ng DT/ 10 µL sterile PBS per injection). Control mice received injections of 10 µL sterile PBS. At days 14 and 28 post-injury, hindlimbs were harvested at the knee and used for functional and biomechanical assessments.

#### Histology and immunofluorescence

Hind paws were harvested (n = 3-5 per timepoint) from uninjured GLAST^Ai9^ and GLAST^Ai9^; Scx^GFP^ mice, and GLAST^Ai9^; Scx^GFP^ at days 3, 7, 10, 14, 21, and 28 days post-injury for paraffin sectioning. Hind paws were fixed in 10% neutral buffered formalin (NBF) for 72 hours at room temperature, decalcified for two weeks, processed, and embedded in paraffin. Three-micron sagittal sections were cut, deparaffinized, and rehydrated. Antigen retrieval (10mM sodium citrate, 0.05% Tween-20, pH 6.0) was performed at 75°C for 3 hours. Slides were blocked in blocking buffer (10% normal donkey serum [NDS; #017-000-121, Jackson ImmunoResearch, West Grove, PA, USA] in PBS + 0.1% Tween-20 [PBST]) for 1 hour at room temperature. For non-conjugated mouse antibodies (*e.g.*, mouse anti-myosin) listed below, 30 μg/mL goat anti mouse IgG Fab fragment (#115-007-003, Jackson ImmunoResearch) was added to the blocking buffer. Slides were then incubated with antibodies to tdTomato (1:500, AB8181, SICGEN, Cantanhede, PT), GFP (1:500, ab290, Abcam, Cambridge, UK), myosin (1:50, ab37484, Abcam), alpha-smooth muscle actin (αSMA-FITC, 1:500, F3777, Sigma Aldrich), CD146 (1:500, ab75769, Abcam), CCN3 (1:100, ab137677, Abcam), Ki67 (1:250, ab16667, Abcam), or CXCL14 (1:750, PA5-28820, Thermo Fisher Scientific, Waltham, MA, USA) in blocking buffer overnight at 4°C. The following secondary antibodies were used (all used at 1:200, all AffiniPure F(ab’)₂ Fragment, Jackson ImmunoResearch): donkey anti-goat Rhodamine-Red-X (RRX) for tdTomato (#705-296-147), donkey anti-rabbit 488 for GFP and Ki67 (#711-546-152), donkey anti-rabbit RRX for CCN3 (#711-296-152), donkey anti-mouse 488 for myosin (#715-546-150), and donkey anti-rabbit Alexa Fluor 647 for CXCL14 (#711-606-152). All secondary antibody incubations were performed for 1 hour at room temperature. Nuclei were counterstained with Hoechst 33342 (NucBlue™ Live ReadyProbes™ Reagent, Thermo Fisher Scientific) and slides were cover slipped (ProLong™ Diamond Antifade Mountant, Thermo Fisher Scientific). Imaging was performed using a VS120 Virtual Slide Microscope (Olympus, Waltham, MA, USA) and images were pseudo-colored in OlyVIA (V3.3, Olympus) for ease of visualization and accessibility.

#### Quantification of cell populations during healing

Fluorescent images were quantified with VIS Image Analysis Software (v.6.7.0.2590, Visiopharm, Hørsholm, DK) as previously described.^60, 62^ Briefly, a region of interest (ROI) was drawn around the scar area (excluding tendon stubs). Individual nuclei were then identified by automatic segmentation and were subsequently assigned a specific identity based on intensity thresholds of each color channel. The total number of cells was determined based on the number of nuclei identified, and the percentage of each cell population is given as a percentage of this total cell number. An n = 3-5 repairs was used for quantification of cell populations (GLAST^Ai9^, GLAST^Ai9^; Scx^GFP^, Scx^GFP^, or double negative) as well as proliferating GLAST^Ai9^ (GLAST^Ai9^; Ki67+) and pro-fibrotic peripheral GLAST^Ai9^ cells (GLAST^Ai9^; CXCL14+) during tendon healing.

#### Human tissue histology

Normal human tendon tissue from uninjured flexor tendons (n= 2) was obtained after informed consent and was approved by the Research Subjects Review Board at the University of Rochester (protocol #54231). In addition, peritendinous scar tissue was obtained from patients (n= 3) undergoing tenolysis procedure. All human samples were processed for routine paraffin histology. Three-micron sagittal sections were cut, deparaffinized, rehydrated, probed with antibodies to GLAST (EAAT1, 1:250, ab416, Abcam) or CCN3 as described above, and visualized using a donkey-anti rabbit RRX secondary antibody. Imaging and pseudo-coloring were performed as described above.

#### Second harmonic generation two-photon microscopy

To visualize the relationship between GLAST^Ai9^ cells and the tendon collagen matrix using second harmonic generation (SHG) imaging, uninjured hind paws from GLAST^Ai9^ mice (n=3) were harvested for frozen sectioning as previously described.^20,65^ Briefly, hind paws were fixed for 24 hours in 10% NBF, decalcified for 5 days in EDTA, and incubated in 30% sucrose for 24 hours to cryo-protect the tissue. 20μm sagittal and axial sections of intact FDL tendons were cut using a cryotape-transfer method^63^ and nuclei were stained with Hoechst 33342 as described above. Z-stack images of endogenous tdTomato expression in GLAST^Ai9^ cells along with the collagen SHG of the tendon body were then acquired in the URMC Multiphoton and Analytical Imaging Center (MAGIC) Core using an Olympus FVMPE-RS system and tunable Insight X3 laser. Images were pseudo-colored and analyzed in Fiji (https://fiji.sc/).^64^ 3D projections and renderings of image stacks were generated using the 3D Project function and 3D Viewer plugin, respectively.

#### *In situ* two-photon microscopy

Hind paws from GLAST^Ai9^ mice were harvested 14 days following the final TMX injection. The skin on the plantar surface of the hind paw was removed, and the superficial-most muscle on the bottom of the hind paw was carefully dissected and removed to expose the FDL tendon. Dissected hind paws were then fixed for 24 hours in 10% NBF at 4°C, protected from light. To visualize the location of GLAST^Ai9^ cells relative to blood vessels on the surface of the tendon, hind paws were stained with antibodies to αSMA which labels vascular smooth muscle cells. Following fixation, hind paws were rinsed, blocked (blocking buffer, 1 hour at room temperature), and incubated with conjugated antibodies to αSMA (αSMA-FITC, 1:500, F3777, Sigma Aldrich) in blocking buffer for 48 hours at 4°C, protected from light. Hind paws were rinsed again, and nuclei were counterstained with Hoechst 33342 for 1 hour at room temperature. For imaging, hind paws were anchored plantar side up in a clear putty (Crazy Aaron’s Thinking Putty®, Liquid Glass®, Crazy Aaron Enterprises, Norristown, PA, USA) and covered with PBS. Z-stack images of the epitenon were acquired in the URMC MAGIC Core and images were analyzed and pseudocolored in Fiji as described above. 3D projections and renderings of image stacks were generated using the 3D Project function and 3D Viewer plugin, respectively.

#### Single cell isolation

##### Mouse

In order to isolate sufficient cells for single cell RNA-sequencing (scRNA-seq), tendons from individual GLAST^Ai9^ mice were pooled for isolation (12x intact tendons, 5x day 7 repairs, 3x day 14 repairs, and 3x day 28 repairs). Intact tendons were harvested from the heel to just proximal to the digit bifurcation. For injured tendons, the entire scar area was carefully dissected away from surrounding tissue and the scar along with approximately 1 mm of tendon stub on either side of the repair was harvested. Tendons were placed into low glucose (1 g/L) Dulbecco’s Modified Eagle Medium (DMEM; 11885084, Gibco, Thermo Fisher Scientific) on ice, and digested in DMEM containing Collagenase Type I (LS004196, Worthington Biochemical, Lakewood, NJ, USA) and Collagenase Type IV (LS004188, Worthington Biochemical) at a final concentration of 5 mg/ml and 1 mg/ml, respectively. Samples were digested at 37°C with rotation for approximately 2 hours for intact tendons and 45 minutes for injured tendons. This protocol was optimized to ensure high quality cells for sequencing. After digestion, the cell suspension was sequentially filtered through 70 μm and 50 μm cell strainers to remove any debris, pelleted at 500xg for 10 minutes at 4°C, rinsed twice in Dulbecco’s Phosphate Buffer Solution (dPBS; 14190136, Gibco), and finally resuspended in 0.5% Bovine Serum Albumin (BSA; Fraction V, 10735078001, Roche, Basel, CH) in dPBS at a concentration of 1,200 cells/μL.

##### Human

Peritendinous scar tissue was obtained from a single individual (34-year-old male with zone 5 lacerations to his flexor pollicis longus and all flexor digitorum profundus/ superficialis tendons) with approval by the Research Subjects Review Board at the University of Rochester (protocol #54231) and following informed patient consent. Eight months following initial repair, the patient underwent a tenolysis procedure to restore digit range of motion. Peritendinous scar tissue was removed during surgery, harvested into low glucose DMEM on ice, and immediately processed for scRNA-seq. Fatty tissue was removed and fibrous scar tissue was minced into small (∼1mm) pieces before being digested in DMEM containing Collagenase Type I and Type IV as described above. Tissue was incubated at 37°C for approximately 1 hour until completely digested before being filtered, rinsed, and resuspended in 0.5% BSA dPBS as described above.

#### Single cell RNA-sequencing

For both species, libraries were prepared and sequenced by the UR Genomics Research Center (GRC). 10,000 cells per sample were loaded into Chromium Controller (10X Genomics, Pleasanton, CA, USA) for single-cell capture. Libraries were then prepared using the Chromium Single Cell 3’ Reagent Kit v3 (#1000075, 10X Genomics). Following library preparation and quality control (QC), libraries were sequenced using the S2 NovaSeq flow cell system (Illumina, San Diego, CA) generating an average of 1.7×10^9^ reads per sample (∼140,000 reads per cell) for mouse and 7.6 x10^6^ reads (∼105,446 per cell) for human. Reads were processed using CellRanger (v6.0.1,10X Genomics) and aligned to either the human reference genome (GRCh38-2020-A) or a modified version of the mouse genome (GRCm38/mm10) containing the mRNA sequence for *tdTomato* from the *ROSA-Ai9* targeting vector (#22799, Addgene, Watertown, MA, USA) to facilitate identification of GLAST^Ai9^ cells in the scRNA-seq datasets.

#### Mouse healing single cell RNA-seq data processing, annotation, and integration

All scRNA-seq analysis was performed using RStudio (v1.4.1717). Datasets from each timepoint were processed individually using the Seurat R package (v4.3.0).^65^ Low quality cells with low feature numbers or mitochondrial gene content greater than 5% were excluded from the dataset, resulting in a total of 6,320 cells for the uninjured sample, 4,470 for the day 7 samples, 4,841 for the day 14 sample, and 4,609 cells for the day 28 sample. Read data for each sample was normalized (normalization.method = “LogNormalize”, scale.factor = 10000) and the top 2000 most variable features were identified (selection.method = “vst”) for use in downstream analysis. Data was scaled, and dimensionality reduction was performed by *RunPCA* using the highly variable features with default parameters. Unsupervised clustering was then performed using the *FindNeighbors* function to compute the shared nearest-neighbors (SNN) for each sample (dims = 1:20) and the *FindClusters* function (resolution = 0.1). Clusters were visualized as Uniform Manifold Approximation and Projection (UMAP) plots in two dimensions. Differentially expressed genes (DEGs) for each cluster were identified using the *FindAllMarkers* function (min.pct = 0.25, logfc.threshold = 0.25). To aid in annotating individual clusters, gene ontology (GO) enrichment analysis was performed with the DAVID Gene Functional Classification Tool (http://david.abcc.ncifcrf.gov; version 6.8)^66^ using the top 100 DEG from each cluster. To trace the fate of GLAST^Ai9^ epitenon cells throughout healing, the uninjured dataset was integrated with datasets from days 7, 14, and 28 post-injury using the *FindIntegrationAnchors* and *IntegrateData* functions (dims = 1:20). Scaled data was used for PCA (npcs = 30) and UMAP dimensional reduction. Cell clusters were visualized (resolution = 0.21) and identified as described above. For both the individual timepoints and the integrated dataset, GLAST^Ai9^ cells were identified as any cell with normalized *tdTomato* expression >0.05. Similarly, Scx+ cells were identified as any cell with normalized *Scx* expression >0.3.

##### Pseudotime and transcription factor enrichment analysis

The tenogenic and pro-fibrotic differentiation trajectories of epitenon cells during healing were modeled using the Monocle3 R package (v.3.1.2.9).^67, 68^ ^71,72^ Epitenon clusters (uninjured, day 7, and day 14) and bridging cell clusters (day 7, day 14, and day 28) were subset from the integrated dataset and converted into a Monocle3 object using the SeuratWrappers R package (v0.3.1). As the bridging cell cluster appears post-injury, no uninjured bridging cell cells were able to be included in the trajectory analysis. The day 28 epitenon cell cluster was excluded from the pseudotime analysis as cells at this timepoint appear to be reverting back to the uninjured epitenon phenotype (**Fig. S6B**) and therefore were not representative of the pro-fibrotic phenotype of interest. Cells were re-clustered (resolution = 0.001) and the functions *learn_graph* and *order_cells* were utilized to order cells in pseudotime and to identify potential differentiation trajectories using the uninjured epitenon cell cluster as the root. The *graph_test* and *find_gene_modules* (resolution = 0.001, k = 13) functions were then used to generate gene modules (groups of genes that vary over pseudotime) by hierarchical cluster analysis. GO enrichment analysis was subsequently performed to determine enriched biological processes in each module. To determine potential transcription factors (TF) that may be involved in regulating either the tenogenic or pro-fibrotic fate of epitenon cells post-injury, transcription factor enrichment analysis was performed using the web-based ChEA3 tool (https://maayanlab.cloud/chea3/).^69^ For each module, the top 5 transcription factors were selected and plotted over pseudotime using the *plot_genes_in_pseudotime* function.

##### Human and mouse scRNA-seq integration

In addition to the peritendinous human scar tissue described above, previously published human peritendinous scar scRNA-seq data from 12 weeks (n = 3) post-surgery was used (SRA: PRJNA975881).^30^ Quality control on these samples was performed as described in the original study.^30^ A total of 42,596 cells derived from human peritendinous scar tissue were used for downstream analyses. Human gene symbols were converted to mouse symbols using the *convert_human_to_mouse_symbols* function in NicheNet (v1.1.1)^70^, retaining only human genes with one-to-one mouse orthologs. Human cells were then mapped to their corresponding mouse clusters using the Seurat *MapQuery* function^71^ setting the integrated mouse healing dataset as the reference and the integrated human peritendinous scar dataset as the query.

###### Functional and biomechanical assessment

The gliding function and biomechanical properties of healing GLAST^Ai9;DTR^ tendons were assessed as previously described^49, 57, 59–61, 72–74^. Briefly, the FDL tendon was dissected at the myotendinous junction, secured between two pieces of tape using superglue, and then loaded incrementally with small weights ranging from 0 to 19 grams. Images were captured after each load and the flexion angle of the metatarsophalangeal (MTP) joint was measured using Image J^64^. The maximum range of motion (ROM) was defined as the MTP flexion angle at 19 grams. Gliding resistance was derived from the changes in MTP flexion angle over the range of applied loads^75^. A concomitant increase in gliding resistance and decrease in MTP flexion angle is indicative of restricted range of motion due to increased peritendinous scar tissue. After gliding/ROM testing, the FDL tendon was released from the tarsal tunnel and the taped proximal end of the tendon and the toes of the hind paw were secured into an Instron 8841 uniaxial testing system (Instron Corporation, Norwood, MA) for single load-to-failure testing at a rate of 30mm/minute.

###### Animal stratification and statistical analysis

Experimental numbers were determined based on previously published work^49, 57, 59–61, 72–74^ and animals were randomly assigned to experimental treatment and outcome groups prior to surgery. Each data point is representative of one mouse. All statistical analyses were performed using Prism (v10.2.1, GraphPad Software, Boston, MA). For all quantitative outcomes, data were assessed for normality using the Shapiro-Wilk test (p<0.05). Differences between experimental groups and controls at each timepoint were determined by unpaired t tests (normally distributed data: GLAST^Ai9;DTR^ ablation: MTP ROM and maximum load, CXCL14 immunofluorescent staining) or Mann-Whitney tests (non-normally distributed data: GLAST^Ai9;DTR^ ablation: gliding and stiffness) using the Šidák-Holm method to correct for multiple comparisons. For all tests, adjusted p values ≤ 0.05 were considered significant. Data are shown as the mean ± the standard deviation and statistical significance is denoted on graphs using the following convention: *p<0.05, **p<0.01, ***p<0.001, ****p<0.0001.

## Acknowledgements

The authors thank the CMSR Histology, Biochemistry and Molecular Imaging (HBMI) Core for technical assistance with preparing samples for histology and the UR Genomics Research Core for assistance with scRNA-sequencing. We would also like to extend our sincere thanks to Yurong Gao, Ph.D. of the Multiphoton and Analytical Imaging Center (MAGIC) Core for her assistance in designing the *in situ* multiphoton imaging experiments.

## Author contributions

Study conception and design: AECN, AEL; Acquisition of data: AECN, EAS, JR, AK, CK; Analysis and interpretation of data: AECN, LB, AEL; Drafting of manuscript: AECN; Revision and approval of manuscript: AECN, LB, EAS, JR, AK, CK, AEL.

---

**Fig. 1 supplement:** (**A**) Representative images demonstrating GLAST^Ai9^ labeled epitenon cells (red) in the flexor carpi ulnaris (FCU), tail, and Achilles tendons of adult mice. (**B**) Representative image demonstrating lack of GLAST^Ai9^ labeling in the patellar tendon epitenon. Very few GLAST ^Ai9^ cells (white arrow) were observed in the epitenon; the majority of GLAST ^Ai9^ cells around the patellar tendon are adipocytes. (**C**) Representative image demonstrating lack of epitenon cell labeling in GLAST^Ai9^ mice without tamoxifen. The flexor digitorum longus (FDL) tendon is outline by yellow dotted lines. Some GLAST ^Ai9^ cells were observed in surrounding tissues (ie. muscle; red cells, white arrows). Quantification of the percentage of each cluster in the uninjured scRNA-seq dataset that was (**D**) GLAST^Ai9^ or (**E**) GLAST^Ai9^; Scx+. (**F**) UMAP plots of markers used to annotate cell clusters in the uninjured scRNA-seq dataset.

**Fig. 3 supplement:** (**A**) Representative images of healing tendons from GLAST^Ai9^; Scx^GFP^ mice at 3, 7, 10, 14, 21, and 28 days post-injury stained for alpha smooth muscle actin (αSMA)+ myofibroblasts (green) and GLAST^Ai9^ (magenta) cells (no Scx^GFP^ staining). White arrows denote GLAST^Ai9^ αSMA+ myofibroblasts in the tendon stubs. No GLAST^Ai9^ αSMA+ were found in the scar area. Scale bars = 200 μm (top), 100 μm (bottom). Sutures are marked by white asterisks. (**B**) (**top**) Representative images of healing tendons from GLAST^Ai9^; Scx^GFP^ mice at 3, 7, 10, and 28 days post-injury demonstrating staining for CD146 (green) and GLAST^Ai9^ (red) cells in the scar area. No GLAST^Ai9^; CD146+ cells were observed in any sample. Scale bars = 50 μm. (**bottom)** Quantification of the mean GLAST^Ai9^ (red), CD146+ (green), GLAST^Ai9^; CD146+ cells (yellow), or double negative cells (blue) in the scar area throughout healing. Error bars denote standard deviation. All images are representative of n = 2-4 mice. In all images, nuclei are counterstained with Hoechst (blue).

**Fig. 4 supplement:** (**A**) Volcano plots of differentially expressed genes (DEG) between GLAST^Ai9^+ and GLAST^Ai9^- in the epitenon and bridging cell clusters. tdTomato was removed as a feature for visualization. (**B**) Comparisons of DEG from different timepoints in the epitenon cluster (volcano plots) and GO term analysis for biological processes enriched at each timepoint (bar graphs). (**C**) Comparisons of DEG from different timepoints in the bridging cell cluster (volcano plots) and GO term analysis for biological processes enriched at each timepoint (bar graphs).

**Fig. 6 supplement:** (**A**) UMAP plot of sub-setted epitenon and bridging cell clusters for pseudotime trajectory analysis, including the day 28 (D28) epitenon cluster. (**B**) Box plot of clusters from (**A**) showing distribution of cells in each cluster over pseudotime. Violin plots of (**C**) tenogenic and (**D**) fibrotic marker expression in real time post-injury.

**Fig. 7 supplement:** (**A**) Experimental overview of depletion and analysis. (**B**) Representative images of uninjured flexor tendons from GLAST^Ai9;DTR^ mice showing GLAST^Ai9^ (magenta) cells prior to depletion (Control), 1 day post depletion, and 5 days post depletion. (C) Quantification of the mean percentage of GLAST^Ai9^ cells present in both the epitenon and tendon before and after depletion. Error bars denote the standard deviation. In all images, nuclei are counterstained with Hoechst (blue). n=3 mice per timepoint. Scale bars = 100 μm.

## References

1. Nichols, A.E.C., Best, K.T., and Loiselle, A.E. The cellular basis of fibrotic tendon healing: challenges and opportunities. Translational Research 209, 156–168 (2019).

2. Pennisi, E. Tending tender tendons. Science (New York, N.Y.) 295, 1011 (2002).

3. Mehrzad, R., Mookerjee, V., Schmidt, S., Jehle, C.C., Kiwanuka, E., and Liu, P.Y. The Economic Impact of Flexor Tendon Lacerations of the Hand in the United States. Ann Plast Surg 83, 419–423 (2019).

4. Strickland, J.W., and Glogovac, S.V. Digital function following flexor tendon repair in Zone II: A comparison of immobilization and controlled passive motion techniques. J Hand Surg Am 5, 537–543 (1980).

5. Quadlbauer, S., Pezzei, C., Jurkowitsch, J., Reb, P., Beer, T., and Leixnering, M. Early Passive Movement in flexor tendon injuries of the hand. Archives of orthopaedic and traumatic surgery. Archiv fur orthopadische und Unfall-Chirurgie 136, 285–293 (2016).

6. Civan, O., Gürsoy, M.K., Cavit, A., Özcanlı, H., and Karalezli, M.N. Tenolysis rate after zone 2 flexor tendon repairs. Jt Dis Relat Surg 31, 281–285 (2020).

7. Garner, W.L., McDonald, J.A., Koo, M., Kuhn, C., 3rd, and Weeks, P.M. Identification of the collagen-producing cells in healing flexor tendons. Plastic and reconstructive surgery 83, 875–879 (1989).

8. Gelberman, R.H., Steinberg, D., Amiel, D., and Akeson, W. Fibroblast chemotaxis after tendon repair. J Hand Surg Am 16, 686–693 (1991).

9. Gelberman, R.H., Vandeberg, J.S., Manske, P.R., and Akeson, W.H. The early stages of flexor tendon healing: a morphologic study of the first fourteen days. J Hand Surg Am 10, 776–784 (1985).

10. Manske, P.R., Lesker, P.A., Gelberman, R.H., and Rucinsky, T.E. Intrinsic restoration of the flexor tendon surface in the nonhuman primate. J Hand Surg Am 10, 632–637 (1985).

11. Khan, U., Edwards, J.C., and McGrouther, D.A. Patterns of cellular activation after tendon injury. J Hand Surg Br 21, 813–820 (1996).

12. Mori, T., Tanaka, K., Buffo, A., Wurst, W., Kühn, R., and Götz, M. Inducible gene deletion in astroglia and radial glia--a valuable tool for functional and lineage analysis. Glia 54, 21–34 (2006).

13. Kan, L., Peng, C.Y., McGuire, T.L., and Kessler, J.A. Glast-expressing progenitor cells contribute to heterotopic ossification. Bone 53, 194–203 (2013).

14. Kan, C., Ding, N., Yang, J., Tan, Z., McGuire, T.L., Lu, H., Zhang, K., Berger, D.M.P., Kessler, J.A., and Kan, L. BMP-dependent, injury-induced stem cell niche as a mechanism of heterotopic ossification. Stem Cell Research & Therapy 10, 14 (2019).

15. Göritz, C., Dias, D.O., Tomilin, N., Barbacid, M., Shupliakov, O., and Frisén, J. A pericyte origin of spinal cord scar tissue. Science (New York, N.Y.) 333, 238–242 (2011).

16. Madisen, L., Zwingman, T.A., Sunkin, S.M., Oh, S.W., Zariwala, H.A., Gu, H., Ng, L.L., Palmiter, R.D., Hawrylycz, M.J., Jones, A.R., Lein, E.S., and Zeng, H. A robust and high-throughput Cre reporting and characterization system for the whole mouse brain. Nat Neurosci 13, 133–140 (2010).

17. Kannus, P. Structure of the tendon connective tissue. Scandinavian journal of medicine & science in sports 10, 312–320 (2000).

18. Yang, S.M., Alvarez, D.D., and Schinder, A.F. Reliable Genetic Labeling of Adult-Born Dentate Granule Cells Using Ascl1^CreERT2^ and Glast^CreERT2^ Murine Lines. The Journal of Neuroscience 35, 15379–15390 (2015).

19. Pryce, B.A., Brent, A.E., Murchison, N.D., Tabin, C.J., and Schweitzer, R. Generation of transgenic tendon reporters, ScxGFP and ScxAP, using regulatory elements of the scleraxis gene. Developmental dynamics : an official publication of the American Association of Anatomists 236, 1677–1682 (2007).

20. Best, K.T., and Loiselle, A.E. Scleraxis lineage cells contribute to organized bridging tissue during tendon healing and identify a subpopulation of resident tendon cells. FASEB journal : official publication of the Federation of American Societies for Experimental Biology 33, 8578–8587 (2019).

21. Muhl, L., Genové, G., Leptidis, S., Liu, J., He, L., Mocci, G., Sun, Y., Gustafsson, S., Buyandelger, B., Chivukula, I.V., Segerstolpe, Å., Raschperger, E., Hansson, E.M., Björkegren, J.L.M., Peng, X.-R., Vanlandewijck, M., Lendahl, U., and Betsholtz, C. Single-cell analysis uncovers fibroblast heterogeneity and criteria for fibroblast and mural cell identification and discrimination. Nature Communications 11, 3953 (2020).

22. Korcari, A., Muscat, S., McGinn, E., Buckley, M.R., and Loiselle, A.E. Depletion of Scleraxis-lineage cells during tendon healing transiently impairs multi-scale restoration of tendon structure during early healing. PloS one 17, e0274227 (2022).

23. Ackerman, J.E., Best, K.T., Muscat, S.N., Pritchett, E.M., Nichols, A.E.C., Wu, C.-L., and Loiselle, A.E. Defining the spatial-molecular map of fibrotic tendon healing and the drivers of Scleraxis-lineage cell fate and function. Cell Reports 41, 111706 (2022).

24. Gumucio, J.P., Schonk, M.M., Kharaz, Y.A., Comerford, E., and Mendias, C.L. Scleraxis is required for the growth of adult tendons in response to mechanical loading. JCI Insight 5 (2020).

25. Lee, C.H., Lee, F.Y., Tarafder, S., Kao, K., Jun, Y., Yang, G., and Mao, J.J. Harnessing endogenous stem/progenitor cells for tendon regeneration. Journal of Clinical Investigation 125, 2690–2701 (2015).

26. Dahlgren, L.A., van der Meulen, M.C., Bertram, J.E., Starrak, G.S., and Nixon, A.J. Insulin-like growth factor-I improves cellular and molecular aspects of healing in a collagenase-induced model of flexor tendinitis. Journal of orthopaedic research : official publication of the Orthopaedic Research Society 20, 910–919 (2002).

27. Kurtz, C.A., Loebig, T.G., Anderson, D.D., DeMeo, P.J., and Campbell, P.G. Insulin-like growth factor I accelerates functional recovery from Achilles tendon injury in a rat model. The American journal of sports medicine 27, 363–369 (1999).

28. Provenzano, P.P., Alejandro-Osorio, A.L., Grorud, K.W., Martinez, D.A., Vailas, A.C., Grindeland, R.E., and Vanderby, R., Jr. Systemic administration of IGF-I enhances healing in collagenous extracellular matrices: evaluation of loaded and unloaded ligaments. BMC Physiol 7, 2 (2007).

29. Lu, J., Chatterjee, M., Schmid, H., Beck, S., and Gawaz, M. CXCL14 as an emerging immune and inflammatory modulator. Journal of Inflammation 13, 1 (2016).

30. Zhang, X., Xiao, Y., Hu, B., Li, Y., Zhang, S., Tian, J., Wang, S., Tao, Z., Zeng, X., Liu, N.N., Li, B., and Liu, S. Multi-omics analysis of human tendon adhesion reveals that ACKR1-regulated macrophage migration is involved in regeneration. Bone Res 12, 27 (2024).

31. Ellis, P.D., Chen, Q., Barker, P.J., Metcalfe, J.C., and Kemp, P.R. Nov gene encodes adhesion factor for vascular smooth muscle cells and is dynamically regulated in response to vascular injury. Arterioscler Thromb Vasc Biol 20, 1912–1919 (2000).

32. Matsushita, Y., Sakamoto, K., Tamamura, Y., Shibata, Y., Minamizato, T., Kihara, T., Ito, M., Katsube, K., Hiraoka, S., Koseki, H., Harada, K., and Yamaguchi, A. CCN3 protein participates in bone regeneration as an inhibitory factor. The Journal of biological chemistry 288, 19973–19985 (2013).

33. Katsuki, Y., Sakamoto, K., Minamizato, T., Makino, H., Umezawa, A., Ikeda, M.A., Perbal, B., Amagasa, T., Yamaguchi, A., and Katsube, K. Inhibitory effect of CT domain of CCN3/NOV on proliferation and differentiation of osteogenic mesenchymal stem cells, Kusa-A1. Biochem Biophys Res Commun 368, 808–814 (2008).

34. Minamizato, T., Sakamoto, K., Liu, T., Kokubo, H., Katsube, K., Perbal, B., Nakamura, S., and Yamaguchi, A. CCN3/NOV inhibits BMP-2-induced osteoblast differentiation by interacting with BMP and Notch signaling pathways. Biochem Biophys Res Commun 354, 567–573 (2007).

35. Lafont, J., Thibout, H., Dubois, C., Laurent, M., and Martinerie, C. NOV/CCN3 induces adhesion of muscle skeletal cells and cooperates with FGF2 and IGF-1 to promote proliferation and survival. Cell Commun Adhes 12, 41–57 (2005).

36. Calhabeu, F., Lafont, J., Le Dreau, G., Laurent, M., Kazazian, C., Schaeffer, L., Martinerie, C., and Dubois, C. NOV/CCN3 impairs muscle cell commitment and differentiation. Experimental cell research 312, 1876–1889 (2006).

37. Marchal, P.O., Kavvadas, P., Abed, A., Kazazian, C., Authier, F., Koseki, H., Hiraoka, S., Boffa, J.J., Martinerie, C., and Chadjichristos, C.E. Reduced NOV/CCN3 Expression Limits Inflammation and Interstitial Renal Fibrosis after Obstructive Nephropathy in Mice. PloS one 10, e0137876 (2015).

38. Abd El Kader, T., Kubota, S., Janune, D., Nishida, T., Hattori, T., Aoyama, E., Perbal, B., Kuboki, T., and Takigawa, M. Anti-fibrotic effect of CCN3 accompanied by altered gene expression profile of the CCN family. J Cell Commun Signal 7, 11–18 (2013).

39. Riser, B.L., Najmabadi, F., Garchow, K., Barnes, J.L., Peterson, D.R., and Sukowski, E.J. Treatment with the matricellular protein CCN3 blocks and/or reverses fibrosis development in obesity with diabetic nephropathy. The American Journal of Pathology 184, 2908–2921 (2014).

40. Lemaire, R., Farina, G., Bayle, J., Dimarzio, M., Pendergrass, S.A., Milano, A., Perbal, B., Whitfield, M.L., and Lafyatis, R. Antagonistic effect of the matricellular signaling protein CCN3 on TGF-beta- and Wnt-mediated fibrillinogenesis in systemic sclerosis and Marfan syndrome. The Journal of investigative dermatology 130, 1514–1523 (2010).

41. Lorda-Diez, C.I., Montero, J.A., Diaz-Mendoza, M.J., Garcia-Porrero, J.A., and Hurle, J.M. Defining the Earliest Transcriptional Steps of Chondrogenic Progenitor Specification during the Formation of the Digits in the Embryonic Limb. PloS one 6, e24546 (2011).

42. Dyment, N.A., Liu, C.-F., Kazemi, N., Aschbacher-Smith, L.E., Kenter, K., Breidenbach, A.P., Shearn, J.T., Wylie, C., Rowe, D.W., and Butler, D.L. The paratenon contributes to scleraxis-expressing cells during patellar tendon healing. PloS one 8, e59944–e59944 (2013).

43. Harvey, T., Flamenco, S., and Fan, C.-M. A Tppp3+Pdgfra+ tendon stem cell population contributes to regeneration and reveals a shared role for PDGF signalling in regeneration and fibrosis. Nature cell biology 21, 1490–1503 (2019).

44. Wang, Y., Zhang, X., Huang, H., Xia, Y., Yao, Y., Mak, A.F.-T., Yung, P.S.-H., Chan, K.-M., Wang, L., Zhang, C., Huang, Y., and Mak, K.K.-L. Osteocalcin expressing cells from tendon sheaths in mice contribute to tendon repair by activating Hedgehog signaling. eLife 6, e30474 (2017).

45. Grinstein, M., Tsai, S.L., Montoro, D., Freedman, B.R., Dingwall, H.L., Villaseñor, S., Zou, K., Sade-Feldman, M., Tanaka, M.J., Mooney, D.J., Capellini, T.D., Rajagopal, J., and Galloway, J.L. A latent Axin2(+)/Scx(+) progenitor pool is the central organizer of tendon healing. NPJ Regen Med 9, 30 (2024).

46. Walia, B., Li, T.M., Crosio, G., Montero, A.M., and Huang, A.H. Axin2-lineage cells contribute to neonatal tendon regeneration. Connective tissue research 63, 530–543 (2022).

47. Howell, K., Chien, C., Bell, R., Laudier, D., Tufa, S.F., Keene, D.R., Andarawis-Puri, N., and Huang, A.H. Novel Model of Tendon Regeneration Reveals Distinct Cell Mechanisms Underlying Regenerative and Fibrotic Tendon Healing. Scientific reports 7, 45238 (2017).

48. De Micheli, A.J., Swanson, J.B., Disser, N.P., Martinez, L.M., Walker, N.R., Oliver, D.J., Cosgrove, B.D., and Mendias, C.L. Single-cell transcriptomic analysis identifies extensive heterogeneity in the cellular composition of mouse Achilles tendons. American journal of physiology. Cell physiology 319, C885–c894 (2020).

49. Ackerman, J.E., Nichols, A.E., Studentsova, V., Best, K.T., Knapp, E., and Loiselle, A.E. Cell non-autonomous functions of S100a4 drive fibrotic tendon healing. eLife 8 (2019).

50. Best, K.T., Mora, K.E., Knapp, E., Buckley, M.R., and Loiselle, A.E. Scleraxis-Lineage Cell Depletion Improves Tendon Healing. bioRxiv (2020).

51. Kendal, A.R., Layton, T., Al-Mossawi, H., Appleton, L., Dakin, S., Brown, R., Loizou, C., Rogers, M., Sharp, R., and Carr, A. Multi-omic single cell analysis resolves novel stromal cell populations in healthy and diseased human tendon. Scientific reports 10, 13939 (2020).

52. Akbar, M., MacDonald, L., Crowe, L.A.N., Carlberg, K., Kurowska-Stolarska, M., Ståhl, P.L., Snelling, S.J.B., McInnes, I.B., and Millar, N.L. Single cell and spatial transcriptomics in human tendon disease indicate dysregulated immune homeostasis. Ann Rheum Dis 80, 1494–1497 (2021).

53. Still, C., Chang, W.-T., Sherman, S.L., Sochacki, K.R., Dragoo, J.L., and Qi, L.S. Single-cell transcriptomic profiling reveals distinct mechanical responses between normal and diseased tendon progenitor cells. Cell Reports Medicine 2, 100343 (2021).

54. Mimpen, J.Y., Ramos-Mucci, L., Paul, C., Kurjan, A., Hulley, P.A., Ikwuanusi, C.T., Cohen, C.J., Gwilym, S.E., Baldwin, M.J., Cribbs, A.P., and Snelling, S.J.B. Single nucleus and spatial transcriptomic profiling of healthy human hamstring tendon. FASEB journal : official publication of the Federation of American Societies for Experimental Biology 38, e23629 (2024).

55. Disser, N.P., Ghahramani, G.C., Swanson, J.B., Wada, S., Chao, M.L., Rodeo, S.A., Oliver, D.J., and Mendias, C.L. Widespread diversity in the transcriptomes of functionally divergent limb tendons. J Physiol 598, 1537–1550 (2020).

56. Steffen, D., Avey, A., Mienaltowski, M.J., and Baar, K. The rat Achilles and patellar tendons have similar increases in mechanical properties but become transcriptionally divergent during postnatal development. The Journal of Physiology 601, 3869–3884 (2023).

57. Best, K.T., Lee, F.K., Knapp, E., Awad, H.A., and Loiselle, A.E. Deletion of NFKB1 enhances canonical NF-κB signaling and increases macrophage and myofibroblast content during tendon healing. Scientific reports 9, 10926 (2019).

58. Ackerman, J.E., and Loiselle, A.E. Murine Flexor Tendon Injury and Repair Surgery. Journal of visualized experiments : JoVE (2016).

59. Best, K.T., Korcari, A., Mora, K.E., Nichols, A.E.C., Muscat, S.N., Knapp, E., Buckley, M.R., and Loiselle, A.E. Scleraxis-lineage cell depletion improves tendon healing and disrupts adult tendon homeostasis. eLife 10 (2021).

60. Muscat, S., Nichols, A.E.C., Gira, E., and Loiselle, A.E. CCR2 is expressed by tendon resident macrophage and T cells, while CCR2 deficiency impairs tendon healing via blunted involvement of tendon-resident and circulating monocytes/macrophages. The FASEB Journal 36, e22607 (2022).

61. Adjei-Sowah, E., Chandrasiri, I., Xiao, B., Liu, Y., Ackerman, J.E., Soto, C., Nichols, A.E.C., Nolan, K., Benoit, D.S.W., and Loiselle, A.E. Development of a nanoparticle-based tendon-targeting drug delivery system to pharmacologically modulate tendon healing. Sci Adv 10, eadn2332 (2024).

62. Nichols, A.E.C., Muscat, S.N., Miller, S.E., Green, L.J., Richards, M.S., and Loiselle, A.E. Impact of isolation method on cellular activation and presence of specific tendon cell subpopulations during in vitro culture. FASEB journal : official publication of the Federation of American Societies for Experimental Biology 35, e21733 (2021).

63. Dyment, N.A., Jiang, X., Chen, L., Hong, S.H., Adams, D.J., Ackert-Bicknell, C., Shin, D.G., and Rowe, D.W. High-Throughput, Multi-Image Cryohistology of Mineralized Tissues. Journal of visualized experiments : JoVE (2016).

64. Schindelin, J., Arganda-Carreras, I., Frise, E., Kaynig, V., Longair, M., Pietzsch, T., Preibisch, S., Rueden, C., Saalfeld, S., Schmid, B., Tinevez, J.-Y., White, D.J., Hartenstein, V., Eliceiri, K., Tomancak, P., and Cardona, A. Fiji: an open-source platform for biological-image analysis. Nature methods 9, 676–682 (2012).

65. Hao, Y., Hao, S., Andersen-Nissen, E., Mauck, W.M., Zheng, S., Butler, A., Lee, M.J., Wilk, A.J., Darby, C., Zager, M., Hoffman, P., Stoeckius, M., Papalexi, E., Mimitou, E.P., Jain, J., Srivastava, A., Stuart, T., Fleming, L.M., Yeung, B., Rogers, A.J., McElrath, J.M., Blish, C.A., Gottardo, R., Smibert, P., and Satija, R. Integrated analysis of multimodal single-cell data. Cell 184, 3573–3587.e3529 (2021).

66. Huang, D.W., Sherman, B.T., Tan, Q., Collins, J.R., Alvord, W.G., Roayaei, J., Stephens, R., Baseler, M.W., Lane, H.C., and Lempicki, R.A. The DAVID Gene Functional Classification Tool: a novel biological module-centric algorithm to functionally analyze large gene lists. Genome biology 8, R183 (2007).

67. Trapnell, C., Cacchiarelli, D., Grimsby, J., Pokharel, P., Li, S., Morse, M., Lennon, N.J., Livak, K.J., Mikkelsen, T.S., and Rinn, J.L. The dynamics and regulators of cell fate decisions are revealed by pseudotemporal ordering of single cells. Nature Biotechnology 32, 381–386 (2014).

68. Qiu, X., Mao, Q., Tang, Y., Wang, L., Chawla, R., Pliner, H.A., and Trapnell, C. Reversed graph embedding resolves complex single-cell trajectories. Nature methods 14, 979–982 (2017).

69. Keenan, A.B., Torre, D., Lachmann, A., Leong, A.K., Wojciechowicz, M.L., Utti, V., Jagodnik, K.M., Kropiwnicki, E., Wang, Z., and Ma’ayan, A. ChEA3: transcription factor enrichment analysis by orthogonal omics integration. Nucleic acids research 47, W212–w224 (2019).

70. Browaeys, R., Saelens, W., and Saeys, Y. NicheNet: modeling intercellular communication by linking ligands to target genes. Nature methods 17, 159–162 (2020).

71. Stuart, T., Butler, A., Hoffman, P., Hafemeister, C., Papalexi, E., Mauck, W.M., III, Hao, Y., Stoeckius, M., Smibert, P., and Satija, R. Comprehensive Integration of Single-Cell Data. Cell 177, 1888–1902.e1821 (2019).

72. Best, K.T., Knapp, E., Ketonis, C., Jonason, J.H., Awad, H.A., and Loiselle, A.E. Tendon Cell Deletion of IKKβ/NF-κB Drives Functionally Deficient Tendon Healing and Altered Cell Survival Signaling In Vivo. bioRxiv (2020).

73. Best, K.T., Studentsova, V., Ackerman, J.E., Nichols, A.E.C., Myers, M., Cobb, J., Knapp, E., Awad, H.A., and Loiselle, A.E. Effects of Tamoxifen on Tendon Homeostasis and Healing: Considerations for the use of Tamoxifen-Inducible Mouse Models. Journal of orthopaedic research : official publication of the Orthopaedic Research Society (2020).

74. Ackerman, J.E., Muscat, S.N., Adjei-Sowah, E., Korcari, A., Nichols, A.E.C., Buckley, M.R., and Loiselle, A.E. Identification of Periostin as a critical niche for myofibroblast dynamics and fibrosis during tendon healing. Matrix biology : journal of the International Society for Matrix Biology 125, 59–72 (2024).

75. Hasslund, S., Jacobson, J.A., Dadali, T., Basile, P., Ulrich-Vinther, M., Soballe, K., Schwarz, E.M., O’Keefe, R.J., Mitten, D.J., and Awad, H.A. Adhesions in a murine flexor tendon graft model: autograft versus allograft reconstruction. Journal of orthopaedic research : official publication of the Orthopaedic Research Society 26, 824–833 (2008).

