## Supplementary figures and images for "Epitenon-derived cells comprise a distinct progenitor population that contributes to both fibrosis and regeneration following acute flexor tendon injury in a spatially-dependent manner"

### Fig. S1

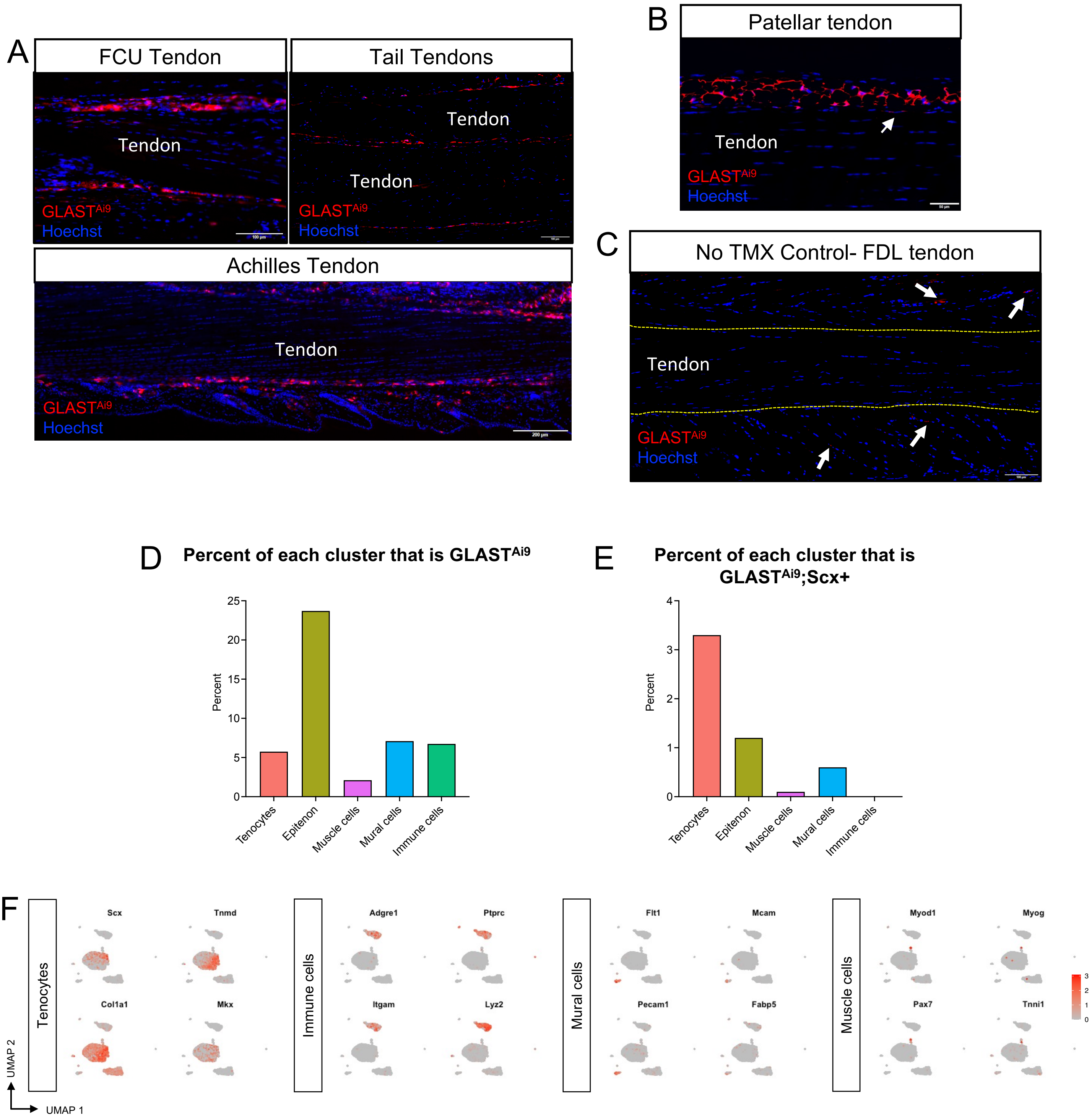

### Fig. S3

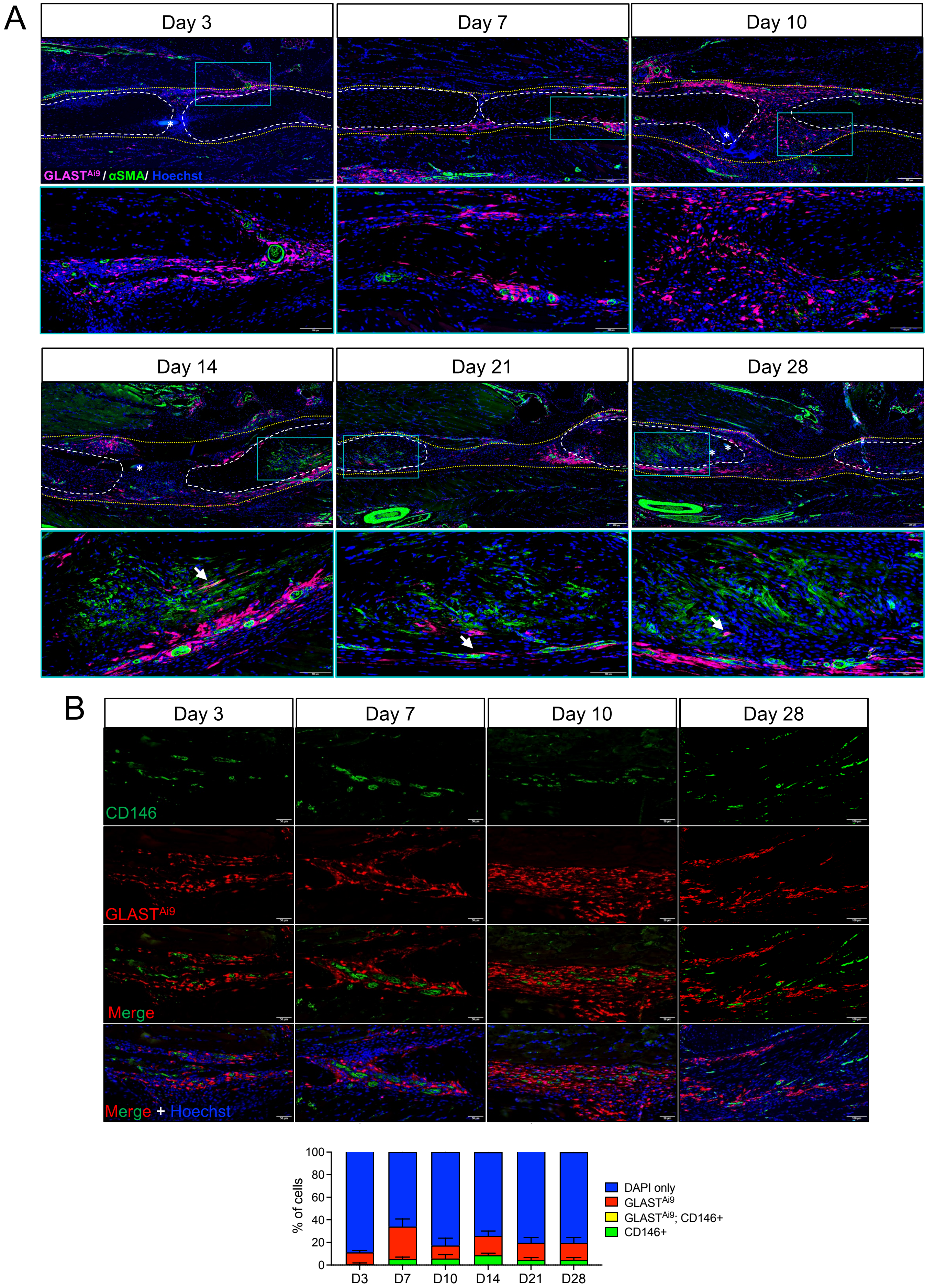

### Fig. S4

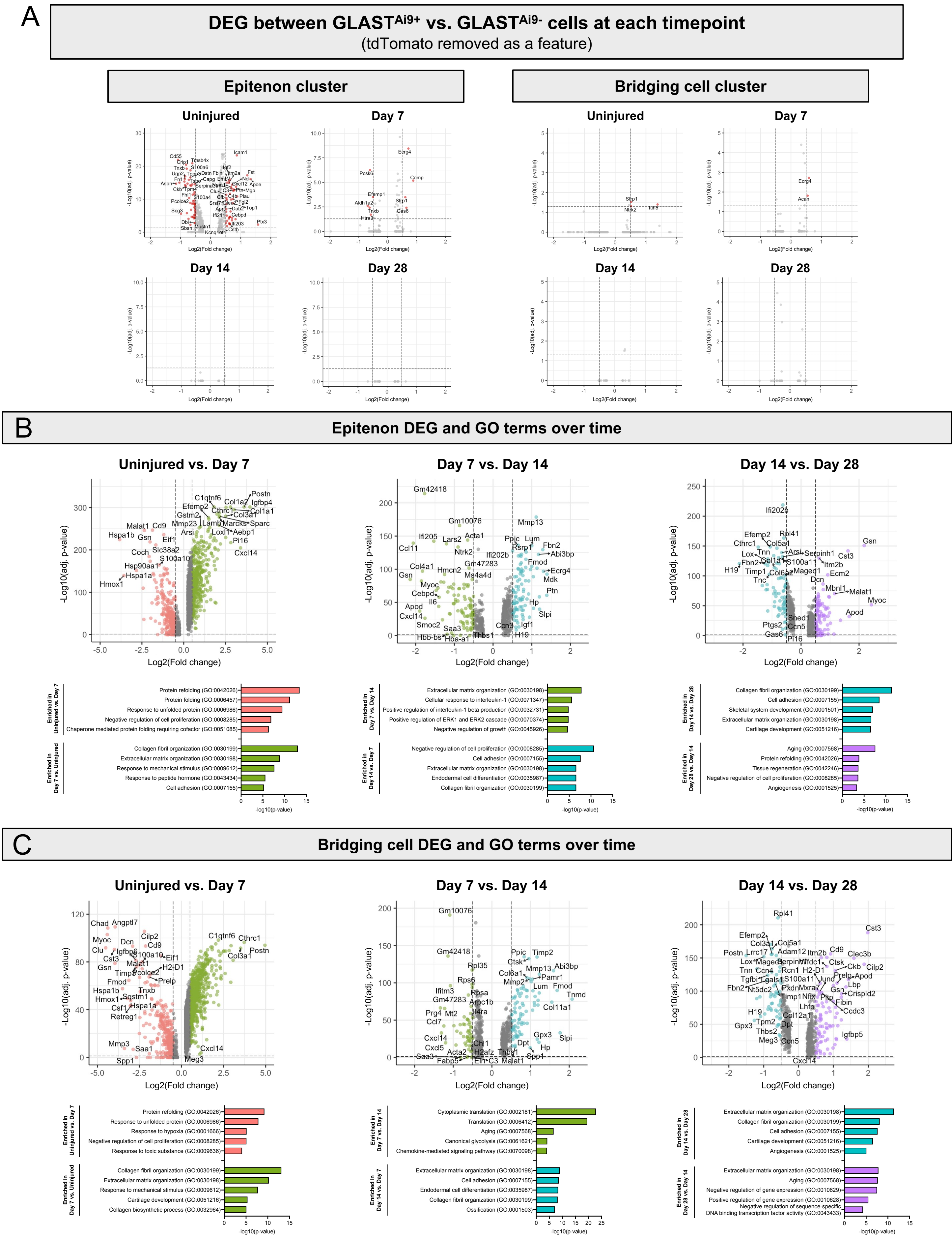

### Fig. S6

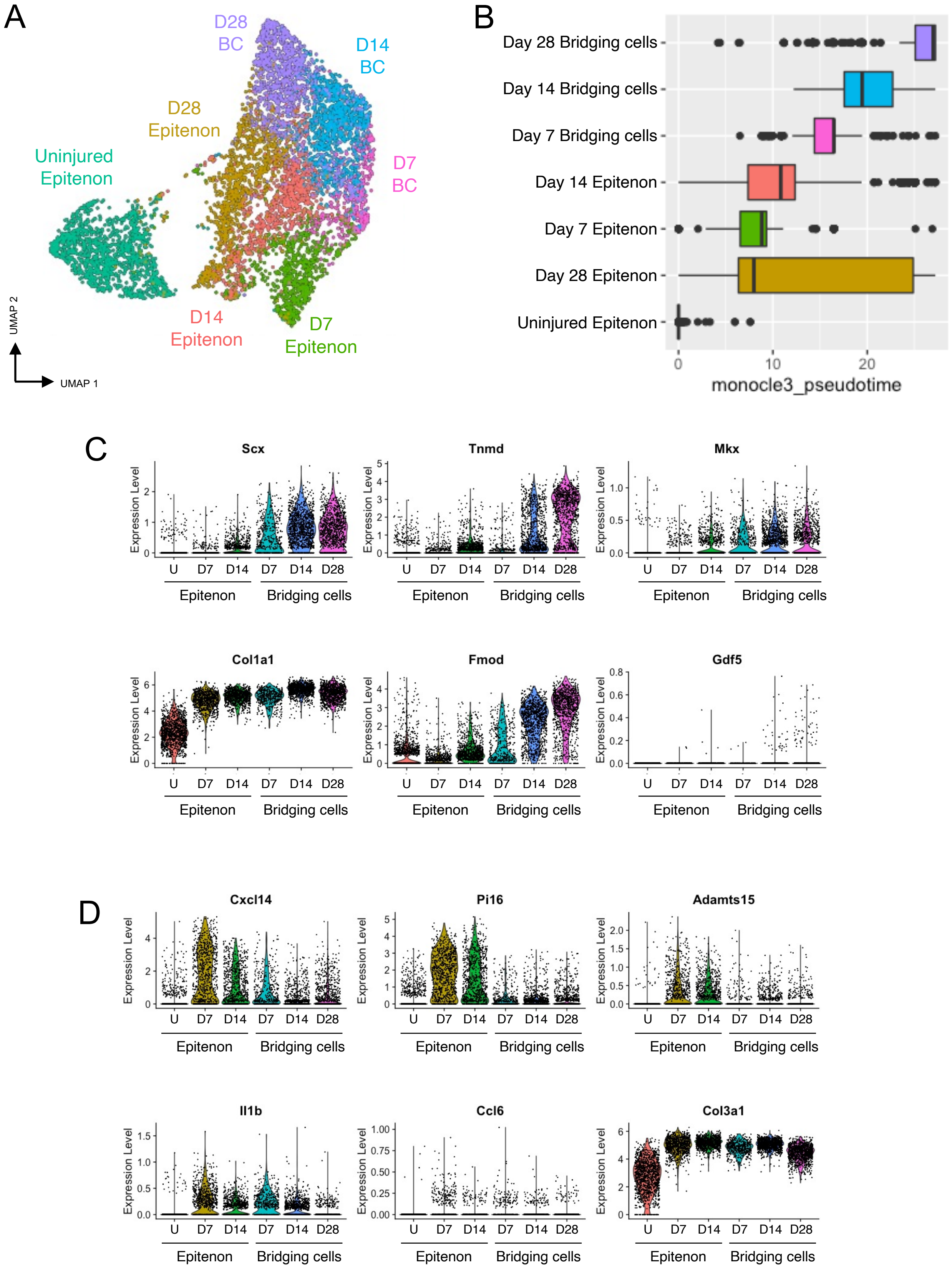

### Fig. S7

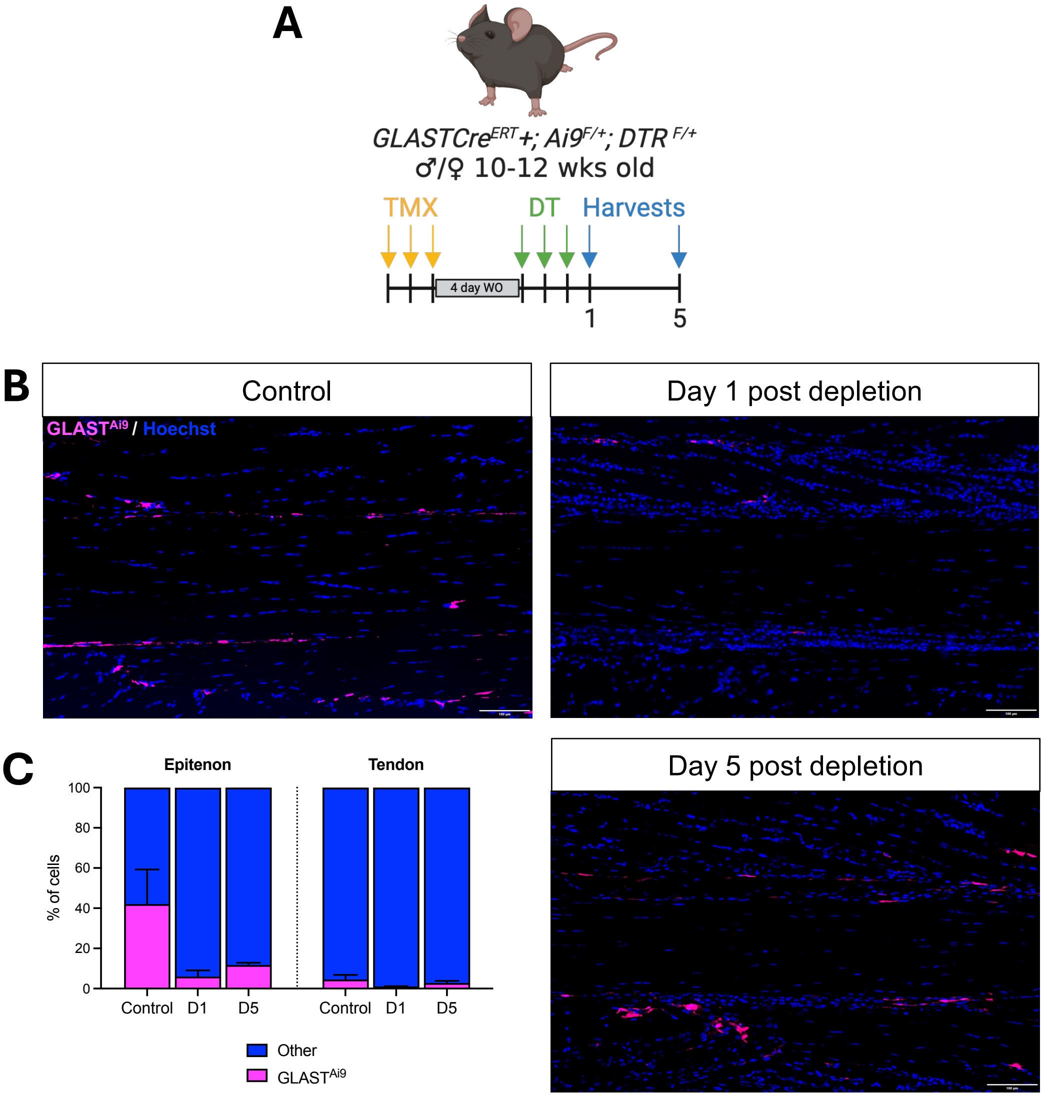

### Fig. S8

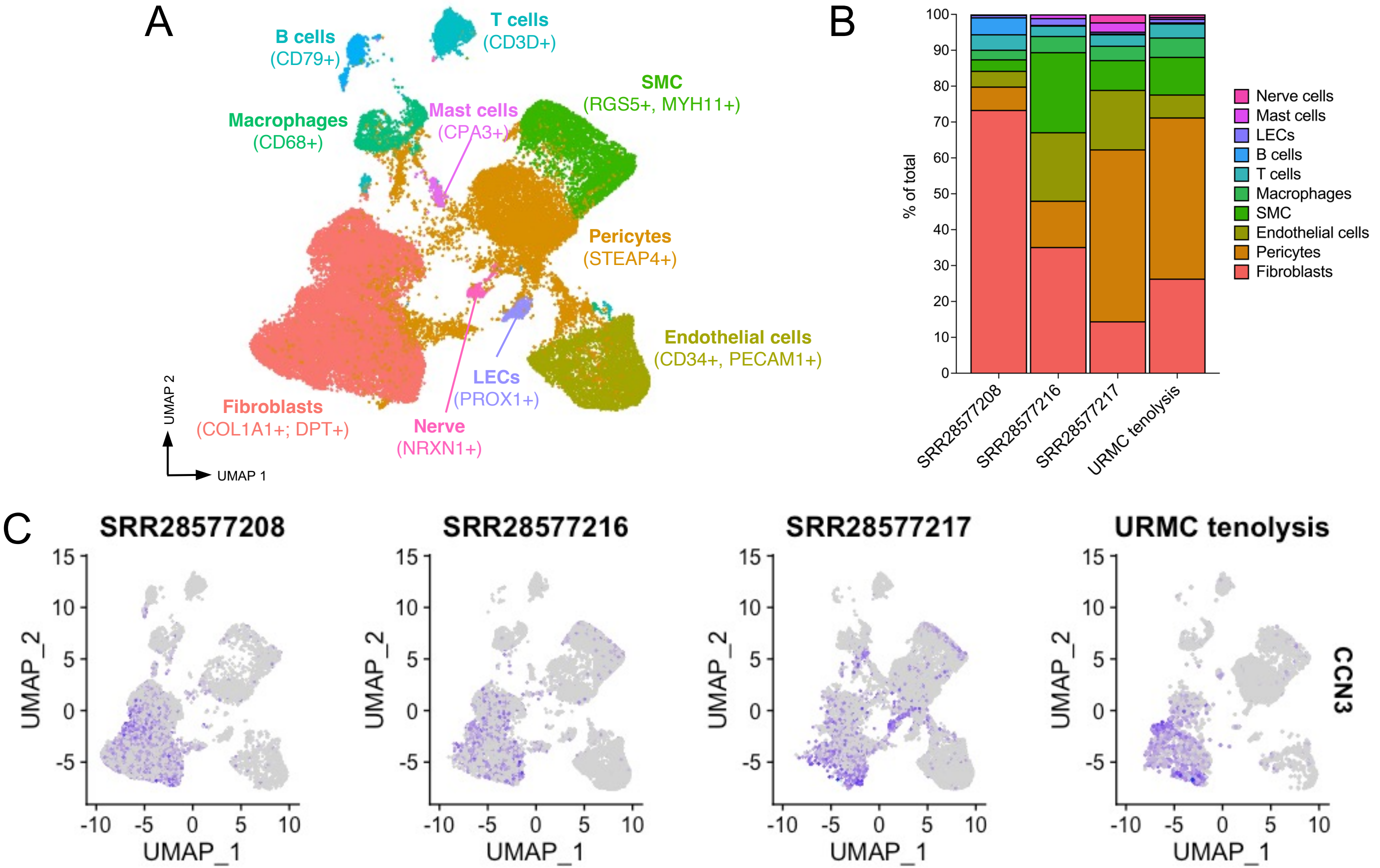
